# N-Cadherin and α-catenin regulate formation of functional tunneling nanotubes

**DOI:** 10.1101/2023.01.10.523392

**Authors:** Anna Pepe, Roberto Notario Manzano, Anna Sartori-Rupp, Christel Brou, Chiara Zurzolo

## Abstract

Cell-to-cell communication it is a fundamental mechanism by which unicellular and multicellular organisms maintain relevant functions as development or homeostasis. Tunneling nanotubes (TNTs) are a type of contact-mediated cell-to-cell communication defined by being membranous structures based on actin that allow the exchange of different cellular material. TNTs have been shown to have unique structural features compared with other cellular protrusions and to contain the cell adhesion molecule N-Cadherin. Here, we investigated the possible role of N-Cadherin and of its primary linker to the actin cytoskeleton, α-Catenin in regulating the formation and transfer function of TNTs. Our data indicate that N-Cadherin through its downstream effector α-Catenin is a major regulator of TNT formation, ultrastructure, as well as of their ability to transfer material to other cells.

## Introduction

Tunneling nanotubes (TNTs) are non-adherent F-actin based membranous structures that form continuous cytoplasmic bridges between cells over distances ranging from several hundred nm up to 100 µm (1). These structures, could be involved both in physiological and pathological conditions (2-6) allowing the transfer of different cargoes (6-11) such as pathogens (12-16) and misfolded proteins between cells (8, 17-22). This ability to transfer a wide variety of cargoes through an open channel connecting two cells is what defines functionally these structures, being therefore unique compared to other cellular protrusions. Despite the large amount of observations supporting the role of TNTs in intercellular communication, the mechanisms of TNT formation and the molecular components and regulators of their structure are still poorly investigated (1). Recently, by developing a correlative light and cryo-electron tomography (ET) workflow, we have shown that TNTs are a unique structure compared to filopodia (23). Indeed, although TNTs appear as single connections by fluorescence microscopy (FM), at nanoscopic level most of them are comprised of a bundle of individual Tunneling Nanotubes (iTNTs) that can contain vesicles and organelles and can be open-ended, thus allowing direct transfer of cellular components. These data also showed long threads coiling around the iTNTs bundles as in the process of holding them together. Of interest, we found the transmembrane protein, N-Cadherin, localized at the attachment point of these threads on the iTNTs membrane, as well as decorating short connections (possibly linkers) between the single tubes (23). N-Cadherin (24) is one of the classical type I cadherins that mediate homophilic cell adhesion dependent on Ca^2+^. N-Cadherin acts through homophilic binding with another N-Cadherin in the opposing cell (25, 26) and interacts with the cadherin-associated molecules p120-Catenin, β-Catenin and α-Catenin (27, 28), the latter being the one that anchors the adhesome to the actin cytoskeleton giving the functionality to this complex (29, 30). α-Catenin also acts as an actin-binding and bundler protein (31) which can also interact with many other actin-binding proteins (32, 33) therefore controlling actin dynamics, as limiting the formation of branched actin filaments (34) or inducing the formation of filopodia by the recruitment of them to PIP3 membranes (35). Based on our and other data showing the presence of members of the cadherins superfamily on TNTs (23, 36-38) we hypothesize that the cadherin-catenin complex could have an important function on the regulation of the TNTs. By combining quantitative assays in living cells with cryo-correlative fluorescent electron microscopy (cryo-CLEM) and tomography here we demonstrate that N-Cadherin is an organizer of the TNT structure and function since the lack of this protein results in disordered and nonfunctional iTNTs. On the other hand, N-Cadherin overexpression increases the stability of these structures and the transfer of vesicles within them. We further demonstrate that α-Catenin its required and is working downstream N-Cadherin in the regulation pathway of TNTs.

## Results

### 1. N-Cadherin interference affects both functionality and ultrastructure of the TNTs

N-Cadherin was previously shown to be present in TNTs in murine neuronal CAD cells (23) as well as in Hela and in urothelial cells (36, 38). To address its role, in this study we used SH-SY5Y human neuronal cells, a relevant model of study for different physiological and pathological neuronal conditions that we had previously characterized for TNT formation (16, 22, 23, 39). Immunofluorescence using anti-N-Cadherin antibody analyzed by confocal microscopy revealed that the transmembrane protein decorated the TNTs formed between these cells (Fig. S1A). Furthermore, immunogold-labeling and cryo-ET showed N-Cadherin localization on the iTNTs membranes and in between iTNTs (Fig. S1B, C), indicating that neuronal cells of different origin (mouse and human) and derivation shared a similar ultrastructural organization of TNTs (23). To investigate a possible role on TNTs, N-Cadherin was knocked-down (KD) in an acute manner by RNA interference (average decrease of 82% compared to RNAi Control) in SH-SY5Y (Fig. S1D).

We quantified the % of TNT-connected cells (see Material and Methods), and found an increase to 56% in RNAi N-Cadherin cells compared to 35% in RNAi Control cells (Fig. 1A, B, C). To test if the increase in TNTs in RNAi N-Cadherin cells correlated to an increase in TNT-mediated vesicle transfer, we performed a transfer assay in a co-culture (40) (Fig. S1E top) (see Material and Methods). After 16h of co-culture, in down-regulated N-Cadherin co-cultures, the percentage of transferred DiD vesicles by a contact-dependent mechanism (17%) was significantly lower compared to the controls (32%) (Fig. 1D-F). In order to rule out any contribution of secretion to the vesicle transfer observed in our co-culture conditions, we performed “secretion tests” in which supernatants from donor RNAi Control and RNAi N-Cadherin cells, were used to challenge acceptor SH-SY5Y cells (Fig. S1E bottom). No significant signal for DiD in the acceptor cells that received the supernatants from the donor cells was found compared to the contact-mediated transfer (Fig. 1F, Fig. S1F), suggesting that the main mechanism of transfer is likely TNT-mediated. In order to understand the mechanisms by which N-Cadherin KD would increase the number of TNT connected cells but decrease TNT-mediated vesicle transfer, we analyzed the structure of TNTs in RNAi N-Cadherin cells by adapting our established cryo-EM and tomography pipeline (16, 23). Similar to mouse neuronal CAD cells (23), TNTs between SH-SY5Y cells were prevalently composed by a bundle of parallel iTNTs (between 2 and 6) (Fig. 1G, H). Strikingly, we observed that in N-Cadherin down-regulated conditions iTNTs did not run parallel (Fig. 1I-J) and braided over each other (Fig.1K). We also observed that compared with the controls, in KD cells there were more tips close-ended (Fig. 1 K, L). These still images could represent iTNT (i) in the process of extending towards other cells, (ii) retracting from opposite cells or (iii) unable to fuse with the opposite cells. To understand whether there was a significative impact of N-Cadherin depletion on the iTNTs morphology, we performed a quantitative analysis of our cryo-EM data to calculate the percentage of close-ended iTNTs vs continuous iTNTs connecting two cells. In control conditions we found that 75% of iTNTs were fully extending between two distant cells, while 25% were interrupted and showed closed tips. In contrast, in RNAi N-Cadherin SH-SY5Y cells, we found a decrease (52%) in fully extended iTNTs (between 2-7 iTNTs per TNT) and an increase of closed tip-iTNTs (48%) (Fig. 1M). It is important to precise that Cryo-ET cannot explore the connecting regions between iTNTs and cell bodies because samples are too thick (>500 nm in thickness), therefore we do not know whether the iTNTs connecting two cell bodies are closed or continuous. However, the images showing disconnected/disjointed and close-ended iTNTs, combined with the decreased in TNT mediated vesicle transfer suggested that in KD cells, TNTs are probably unable to engage and/or fuse with the cell body of the opposing cell, consequently preventing transfer of material.

**Fig. 1.**
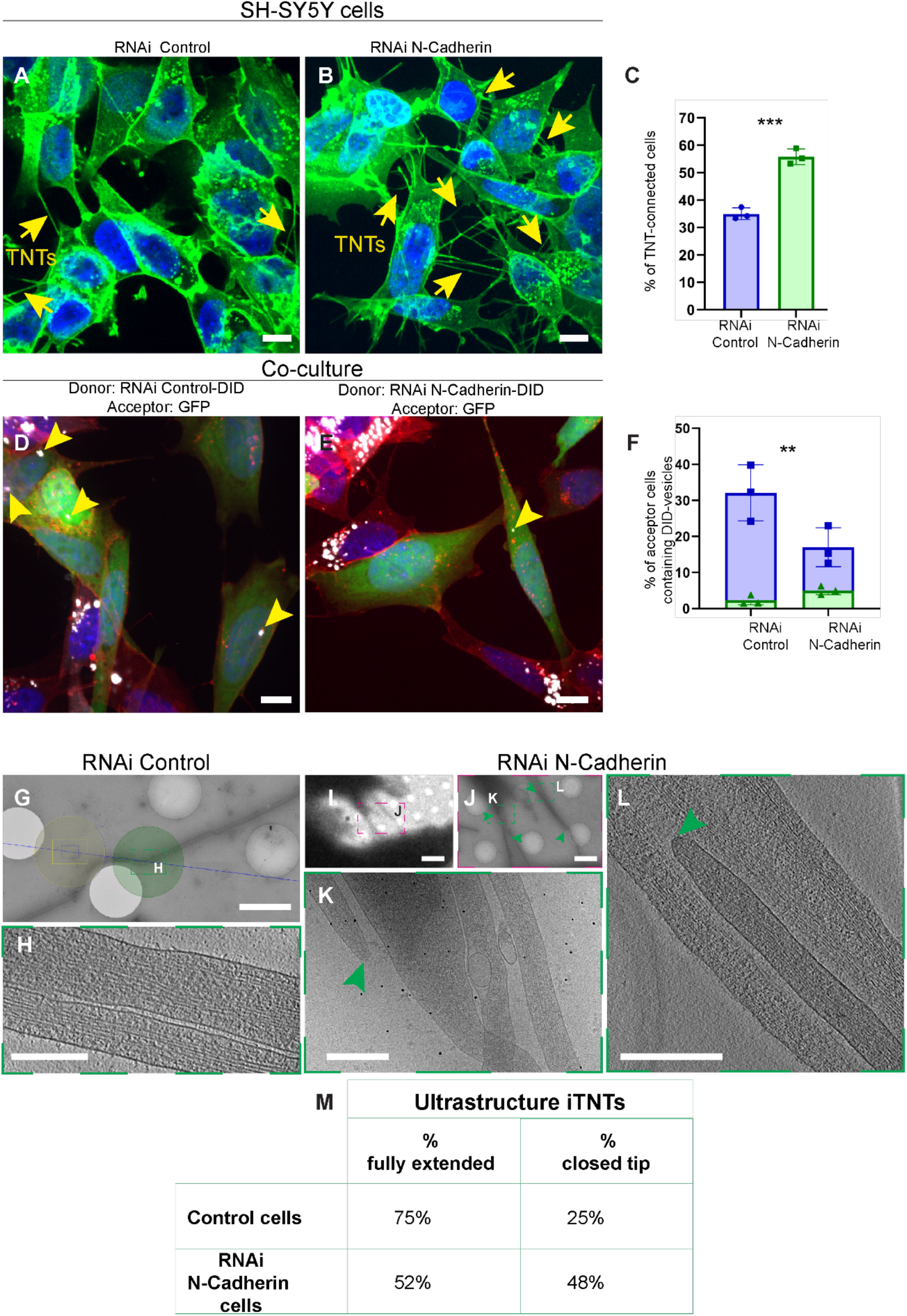
N-Cadherin interference impacts the functionality and ultrastructure of the TNTs between SH-SY5Y cells. (A, B). Confocal micrograph showing (A) TNTs between RNAi Control and (B) TNTs between RNAi N-Cadherin cells. Cells stained with WGA-488 (green) and DAPI (blue) for the nuclei. The yellow arrows indicate the TNTs. (C). Graph showing the percentage of TNT-connected cells transfected with RNAi Control non-targeting (35% ± 2.17) and RNAi N-Cadherin (55.8% ± 2.85), (***p<0.0001 for RNAi Control versus RNAi N-Cadherin for N=3). (D, E). Representative confocal images showing 24h co-culture between (D) RNAi Control with DiD-labelled vesicles (donor) and GFP cells (acceptor), (E) RNAi N-Cadherin challenged with DiD-labelled vesicles (donor) and GFP cells (acceptor). Cellular membranes were labelled with WGA-546 (red), nuclei were stained with DAPI (blue). The yellow arrowheads indicate DiD-labelled vesicles detected in the cytoplasm of acceptor cells. (F). Graph showing the percentage of acceptor cells containing DiD-labelled vesicles from the co-cultures in cells transfected with RNAi Control (32.12%± 7.77 for contact-mediated transfer in blue; 2.31%± 1.31 for transfer by secretion in green) or RNAi N-Cadherin (17%± 5.4 for contact-mediated transfer in blue; 5.03%± 1.18 for transfer by secretion in green) (**p=0.0013 for RNAi Control versus RNAi N-Cadherin for N=3). (G). Cryo-EM intermedia micrograph showing TNT-connected RNAi Control cells. (H). High-magnification cryo-tomography slices corresponding to the green dashed squares in (G) showing full extended iTNTs. (I-L). TNT-connected RNAi N-Cadherin cells acquired by cryo-EM (I) low (J) and intermediate magnification. (K). High-magnification cryo-EM slices showing the iTNT in the green dashed square in (J). (L). High-magnification cryo-tomography slices corresponding to the green dashed squares in (J). (M). Table showing the percentage of full extended iTNTs and closed tip in RNAi Control cells and RNAi N-Cadherin cells. Scale bars in (A, B, D, E) 20 µm, (G, J) 2 µm, (I) 10 µm, (H, K, L) 200nm.

### 2. Overexpression of N-Cadherin promotes TNT-mediated transfer and impacts TNT ultrastructure

To further investigate the effect of N-Cadherin on TNTs we produced an SH-SY5Y cell line stably overexpressing (OE) GFP N-Cadherin, where ectopically expressed GFP N-Cadherin showed a similar cellular distribution compared to the endogenous protein (Fig. S1G). We found that GFP-N-Cadherin cells had a decreased percentage of TNT-connected cells (10%) compared to control cells transfected only with GFP (33%) (Fig. 2A-C). However, these cells exhibited an epithelial-like morphology, where cells tended to stay close to each other in clusters (Fig. S1G), even when we seeded in low concentration to favor a sparse distribution, optimal for the study of TNTs. Therefore, one possibility is that the reduction in TNTs resulted from the fact that TNTs formed by these cells were hidden by the cell bodies in the clusters. To investigate this possibility, we treated the cells with a short pulse of trypsin to the culture to separate the cell bodies and observe the TNTs formed between them (41). After trypsin addition, the % of TNT connected cells increases conspicuously compared to *wild-type* conditions (around 2.5 times more TNTs when we treat the cells with trypsin than in *wild-type* conditions) (Fig. S1H). However, also in this condition GFP N-Cadherin cells formed significantly less TNTs than control cells (respectively 70% compared to 85%) (Fig. S1H-J), confirming that N-Cadherin OE resulted in a reduction of TNT connected cells (Fig. 2A-C). Next, to assess TNT’s functionality we performed a co-culture where GFP N-Cadherin cells loaded with DiD were used as donors and SH-SY5Y mCherry cells as acceptors. Contact-mediated DiD labelled vesicles transfer was significantly higher (40 %) compared to control co-cultures (25 %) where GFP N-Cadherin cells were used as donors (Fig. 2D-F). Importantly, negligible transfer of DiD labelled vesicles by secretion was observed both in control and in GFP N-Cadherin overexpression conditions (Fig. 2F). Thus, the decrease in TNT formation by ectopic expression of N-Cadherin was accompanied by an increase in their transfer function, opposite compared to N-Cadherin downregulation. By cryo-TEM we found that in cells overexpressing N-Cadherin the percentage of TNTs formed by a single tube (65%) was much higher than the % of TNTs formed by an iTNTs bundle (35%) (Fig. 2G-I). This was almost the opposite compared to control SH-SY5Y cells where 40% of TNTs were formed by a single tube while 60% cells were formed by 2 or more iTNTs (Fig. 2I). Interestingly in RNAi N-Cadherin cells the % of single tube TNTs decreased even more (26,3%) and the majority of TNTs (73,7%) were formed by 2 or more iTNTs (Fig. 2I). In addition, 3D Cryo-TEM images revealed that contrary to KD cells (Fig. 1I-L, Fig. S2A-B) in GFP N-Cadherin cells the iTNTs were straight and ran mostly parallel towards the opposite cell (Fig. S2C-G). Quantitative analysis revealed a tendency to have more fully extended iTNTs (58%) and less close-ended tips (42%) (Fig. S2H) compared to the KD cells (52% and 48% respectively) (Fig. 1M). Of interest in our tomograms, we could also discern thin structures connecting the plasma membrane of two iTNTs (Fig. S2G, Movie S1). All these changes in the architecture of TNTs in cells overexpressing N-Cadherin could contribute to the increased transfer that we observed in these conditions (Fig. 2F).

**Fig. 2.**
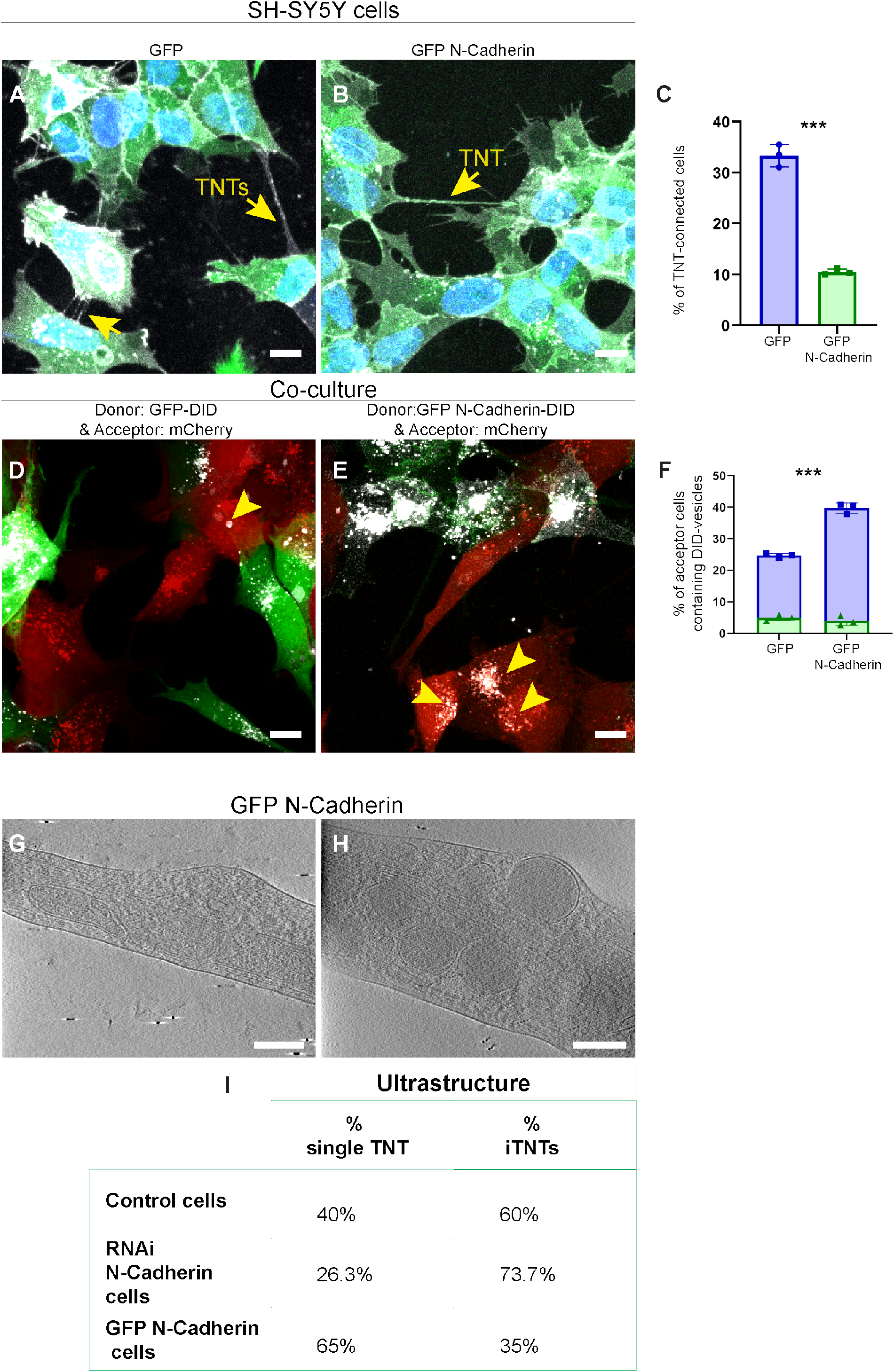
N-Cadherin overexpression impacts the functionality and ultrastructure of the TNTs between SH-SY5Y cells. (A, B). Confocal micrograph showing (A) TNTs between GFP expressing cells, (B) TNTs between GFP N-Cadherin cells. Cells stained with WGA-647 (gray) and DAPI (blue) for the nuclei. The yellow arrows indicate the TNTs. (C). Graph showing the percentage of TNT-connected cells in GFP expressing cells (33.4% ± 2.22) and GFP N-Cadherin (10.5%± 0.60), (***p<0.0001 for GFP versus GFP N-Cadherin for N=3). (D, E). Representative confocal images showing 24h co-culture between (D) GFP with DiD-labelled vesicles (donor) and mCherry cells (acceptor), (E) GFP N-Cadherin challenged with DiD-labelled vesicles (donor) and mCherry cells (acceptor). The yellow arrowheads indicate DiD-labelled vesicles detected in the cytoplasm of acceptor cells. (F). Graph showing the percentage of acceptor cells containing DiD-labelled vesicles from the co-cultures in GFP control cells (24.78%± 0.63 for contact-mediated transfer in blue; 4.97%± 0.93 for transfer by secretion in green) against GFP N-Cadherin cells (39.71%± 1.62 for contact-mediated transfer in blue; 3.95%± 1.48 for transfer by secretion in green). (***p<0.0001 for GFP versus GFP N-Cadherin for N=3). (G, H). High-magnification cryo-tomography slices showing single TNT-connected GFP N-Cadherin cells. (L). Table showing the percentage of single TNTs and iTNTs in Control, RNAi N-Cadherin and GFP N-Cadherin cells. Scale bars in (A, B, D, E) 10 µm, (G, H) 100nm.

### 3. N-Cadherin enhances the stability of the TNTs

One possible explanation for the decrease or increase in vesicle transfer, respectively in N-cadherin KD and OE conditions, is that N-cadherin would regulate formation of functional TNT by favoring a stable attachment of the TNT tip with the opposing cell or by stabilizing the iTNT bundles. To test our hypothesis, we measured by live imaging the lifetime of already formed TNTs in N-Cadherin KD and OE cells compared to respective control conditions (see Material and Methods) (see example on Fig. S3A and Movie S2). In RNAi N-Cadherin cells we observed a clear decrease tendency in the duration of TNTs compared to the respective RNAi Control (18.5 minutes vs 22.5 minutes) (Fig. S3B and Movies S3, Movies S4 respectively). On the other hand, the duration of TNTs in GFP N-Cadherin cells was significative increased compared to its control (GFP expressing cell), (34.5 minutes vs 20 min) (Fig. S3C and Movie S5). Thus, although in N-Cadherin OE there are less TNTs, these are fully formed and more stable, explaining the higher vesicle transfer. On the contrary in N-Cadherin KD there are more TNTs, but these are more disorganized, have more close-ended tips and they seemed to be less stable, thus resulting in lower vesicle transfer.

### 4. N-Cadherin and *α*-Catenin cooperate in TNT regulation

N-Cadherin may affect TNT stability and facilitate vesicle transfer by providing an adhesion complex to bridge connected cells or maintaining the iTNT bundle. To this end we decided to investigate the possible role of α-Catenin, which forms a complex with N-Cadherin mediating the interaction with the actin cytoskeleton, providing integrity of the complex and strengthening adhesion (26). We assessed first the distribution of α-Catenin in our cell model (Fig. S4A). α-Catenin is endogenously expressed by SH-SY5Y cells and is localized at the plasma membrane, in the cytoplasm and on TNTs (Fig. S4A), where it largely co-localizes with N-Cadherin (Fig. S4A). To investigate the involvement in TNT regulation, α-Catenin was KD in an acute manner by RNAi (average decrease of 85% compared to RNAi Control) (Fig. S4B), and TNTs were imaged and quantified. Similar to N-Cadherin KD, the percentage of TNT-connected cells increased to 48% in the α-Catenin depleted cells compared to 29% in RNAi Control cells (Fig. 3A-C). Furthermore, as for N-Cadherin KD, we found that in KD α-Catenin co-cultures the percentage of transferred DiD labelled vesicles (18%) was significantly lower compared to control cells (29%) with insignificant transfer by secretion in both conditions (Fig. 3D-F). On the other hand, upon α-Catenin OE (mEmerald α-Catenin), the percentage of cells connected by TNTs was significatively reduced to around 14% compared to 29% in control conditions (Fig. 3G-I). Despite this reduction, the contact-dependent vesicle transfer showed a significant increase, from 25% in control to 33% of acceptor cells containing transferred vesicles in mEmerald α-Catenin cells with a minimal contribution of the transfer being by secretion (Fig. 3J-L). Once more, these results were in line with the results we obtained by overexpressing N-Cadherin (Fig. 2A&C). To understand if the overlapping functional consequences of N-Cadherin and α-Catenin OE/KD corresponded to similar effects on the ultrastructure and organization of TNTs we analysed by cryo-TEM the morphology of TNTs in cells where α-Catenin was up or down-regulated. Similar to N-Cadherin, in SH-SY5Y cells down-regulated for α-Catenin, iTNTs did not run parallel (Fig. 4A-F), were braided over each other (Fig. 4A, C, D, F) and most of iTNTs were close-ended (Fig. 4A-F), while in mEmerald α-Catenin cells were running mostly parallel (Fig. 4H-L). In addition, 34,4% of iTNTs of RNAi α-Catenin cells were fully extended, while 65% showed closed tip. In contrast, in mEmerald α-Catenin cells, we found a decrease (28%) of closed iTNTs and an increase of fully extended iTNTs (72%) (Fig. 4M).

**Fig. 3.**
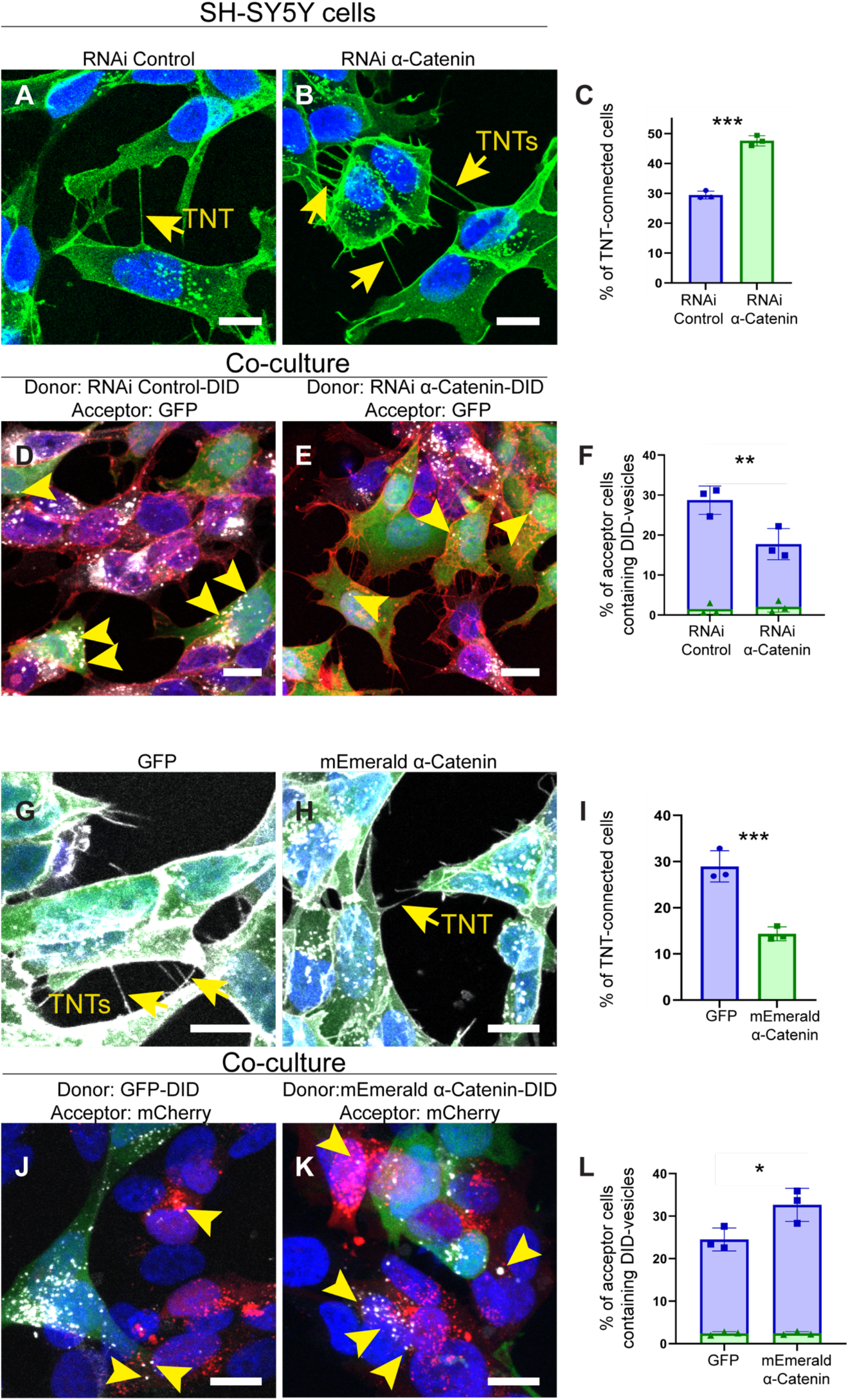
α-Catenin interference and overexpression impact the formation and functionality of the TNTs between SH-SY5Y cells. (A, B). Confocal micrographs showing TNTs between (A) RNAi Control cells, (B) RNAi α-Catenin cells. Cells stained with WGA-488 (green) and DAPI (blue) for the nuclei. The yellow arrows indicate the TNTs. (C). Graph showing the percentage of TNT-connected cells transfected with RNAi Control non-targeting (29.4% ± 1.31) and RNAi α-Catenin (47.6% ± 1.71), (***p=0.0001 for RNAi Control versus RNAi α-Catenin for N=3). (D, E). Representative confocal images showing 24h co-culture between (D) RNAi Control challenged with DiD-labelled vesicles (donor) and GFP cells (acceptor), (E) RNAi α-Catenin with DiD-labelled vesicles (donor) and GFP cells (acceptor). Cellular membranes were labelled with WGA-546 (red), nuclei were stained with DAPI (blue). The yellow arrowheads indicate DiD-labelled vesicles detected in the cytoplasm of acceptor cells. (F). Graph showing the percentage of acceptor cells containing DiD-labelled vesicles from the co-cultures in cells transfected with RNAi Control (28.74%± 3.55 for contact-mediated transfer in blue; 1.5%± 1.33 for transfer by secretion in green) or RNAi α-Catenin (17.75%± 3.91 for contact-mediated transfer in blue; 2.08%± 1.35 for transfer by secretion in green). (**p=0.001 for RNAi Control versus RNAi α-Catenin for N=3). (G, H). Confocal micrograph showing TNTs between (G) GFP cells, (H) mEmerald α-Catenin cells. Cells stained with WGA-647 (grey) and DAPI (blue) for the nuclei. Yellow arrows indicate TNTs connecting two cells. (I). Graph showing the percentage of TNT-connected cells transfected with GFP (29% ± 3.38) and mEmerald α-Catenin (14.3%± 1.49), (***p<0.0001 for GFP versus mEmerald α-Catenin for N=3). Cells stained with WGA-647 (gray) and DAPI (blue) for the nuclei. (J, K). Representative confocal images showing 24h co-culture between (J) GFP with DiD-labelled vesicles (donor) and mCherry cells (acceptor), (K) mEmerald α-Catenin challenged with DiD-labelled vesicles (donor) and Cherry cells (acceptor). The yellow arrowheads indicate DiD-labelled vesicles detected in the cytoplasm of acceptor cells. (L). Graph showing the percentage of acceptor cells containing DiD-labelled vesicles from the co-cultures in GFP control cells (24.52%± 2.70 for contact-mediated transfer in blue; 2.42%± 0.42 for transfer by secretion in green) against mEmerald α-Catenin cells (32.64%± 3.90 for contact-mediated transfer in blue; 2.45%± 0.39 for transfer by secretion in green). (*p=0.0324 for GFP versus mEmerald α-Catenin for N=3). Scale bars, 10 µm.

**Fig. 4.**
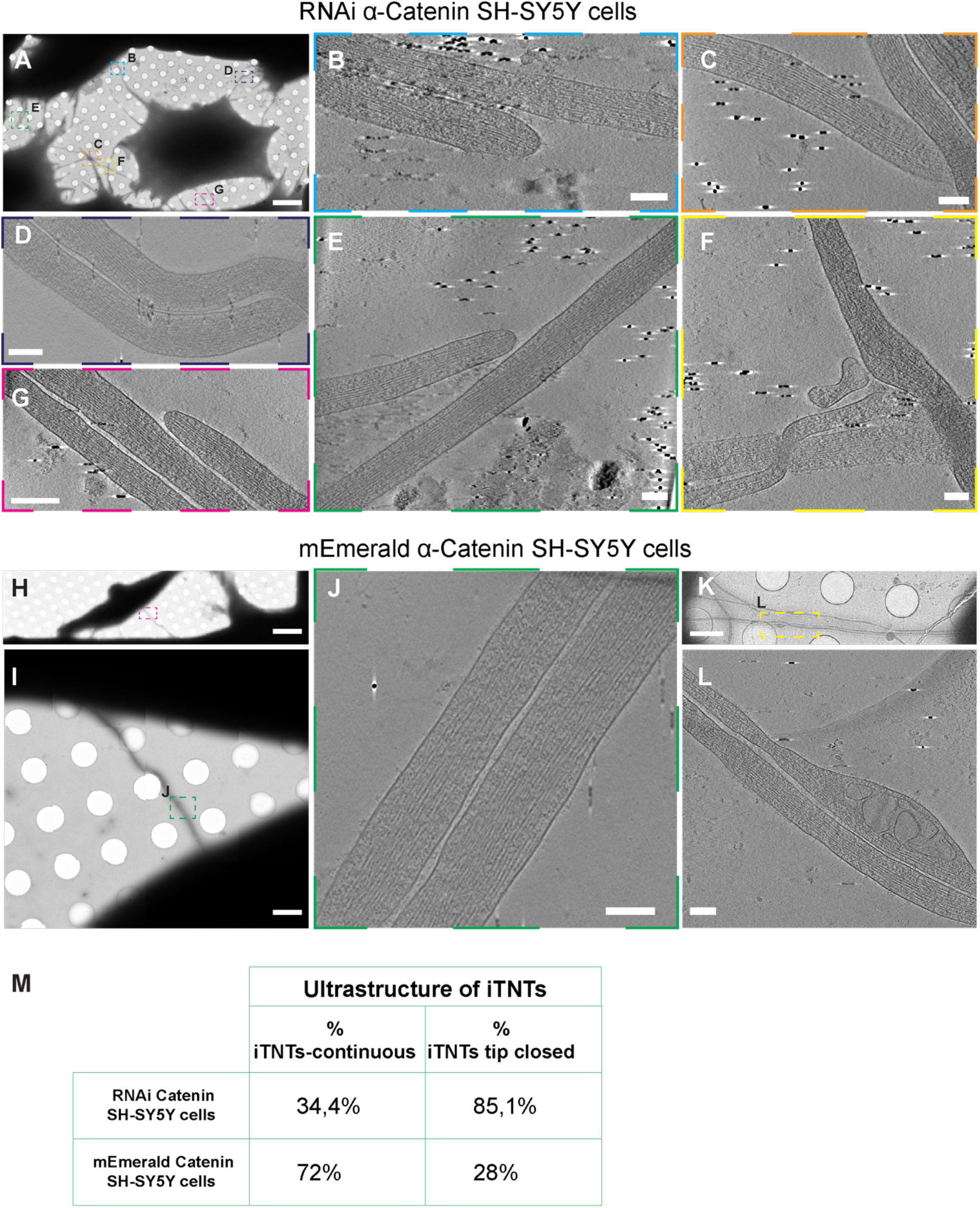
Cryo-EM on TNTs formed between SH-SY5Y in which α-Catenin was up or down-regulated. (A-F). Cryo-EM grids were prepared using RNAi α-Catenin cells (A) Low magnification of cryo-EM micrograph showing TNTs connecting RNAi α-Catenin cells. (B-F) High-magnification cryo-tomography slices of the dashed square in (A) showing iTNTs with closed tip (B, E, G) and iTNTs not running parallel and braided over each other (C, D, E, F). (H-L). Cryo-EM grids were prepared using mEmerald α-Catenin cells. Low (H) and intermedia magnification of cryo-EM micrograph showing TNT connecting mEmerald α-Catenin cells. High-magnification cryo-tomography slices of the green dashed square in (I) showing parallel and fully extended iTNTs. (K) Intermediate magnification of cryo-EM micrograph showing TNT connecting mEmerald α-Catenin SH-SY5Y cells. (L) High-magnification cryo-tomography slices of the yellow dashed square in (K) showing parallel and fully extended iTNTs with vesicles inside. Scale bars (A, H) 10 µm; (I, K) 2µm; (B-F, J, L) 100nm.

### 5. N-Cadherin regulation of TNTs requires *α*-Catenin

To investigate whether the effect of N-Cadherin was mediated by the downstream activity of α-Catenin, we decided to KD α-Catenin in cells overexpressing N-Cadherin. RNAi mediated KD of α-Catenin in GFP N-Cadherin cells and in control cells transfected with mCherry lead to a significative reduction in the levels of this protein compared to the RNAi Control (74% of reduction in GFP N-Cadherin/RNAi α-Catenin cells and 80% in mCherry/RNAi α-Catenin cells) (Fig. S4C). Importantly, N-Cadherin levels and subcellular location were not significatively affected by the downregulation of α-Catenin in GFP N-Cadherin cells (Fig. S4D). We found that α-Catenin KD in GFP N-Cadherin cells led to a significant increase in TNT connected cells compared to RNAi Control cells (from around 8% to almost 19%) (Fig. 5A-C). Furthermore, DiD-vesicle transfer assay in coculture revealed that GFP N-Cadherin/RNAi α-Catenin cells transferred significantly less vesicles (24%) compared to RNAi Control conditions (44%) by a contact-mediated mechanism (Fig. 5D-F), again with a low transfer by secretion compared to the total transfer. These data showed that KD of α-Catenin overcomes almost completely the effect of N-Cadherin overexpression, suggesting that α-Catenin acts downstream N-Cadherin in the regulation of TNTs.

**Fig. 5.**
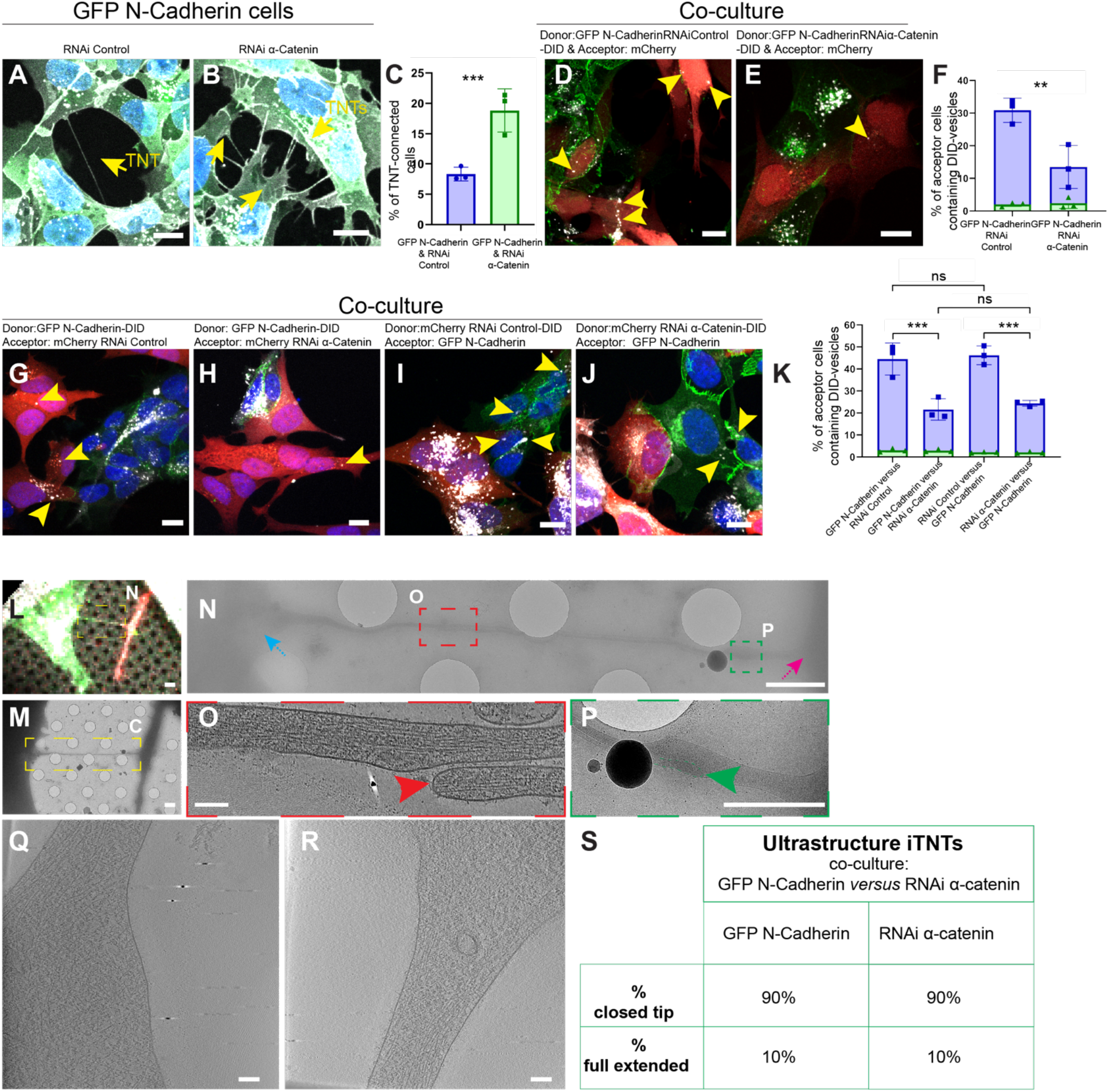
Formation, functionality and ultrastructure of TNTs between N-Cadherin OE and α-Catenin KD cells in co-culture (A, B). Confocal micrograph showing (A) TNTs between GFP N-Cadherin cells with RNAi Control and (B) TNTs between GFP N-Cadherin cells with RNAi α-Catenin. Cells stained with WGA-647 (gray) and DAPI (blue) for the nuclei. The yellow arrows indicate the TNTs connected cells. (C). Graph showing the percentage of TNT-connected GFP N-Cadherin cells transfected with RNAi Control (8.32% ± 1.15) and RNAi α-Catenin (18.8% ± 3.54), (***p<0.0001 for RNAi Control versus RNAi α-Catenin for N=3). (D, E). Representative confocal images after a 24h co-culture between (D) GFP N-Cadherin RNAi Control with DiD-labelled vesicles (donor) and mCherry cells (acceptor), (E) GFP N-Cadherin RNAi α-Catenin with DiD-labelled vesicles (donor) and mCherry cells (acceptor). The yellow arrowheads indicate DiD-labelled vesicles detected in the cytoplasm of acceptor cells. (F). Graph showing the percentage of acceptor cells containing DiD-labelled vesicles from the cocultures in GFP N-Cadherin cells transfected with RNAi Control (43.93%± 2.59 for contact-mediated transfer in blue; 2.24%± 0.22 for transfer by secretion in green) or RNAi α-Catenin (23.53%± 5.23 for contact-mediated transfer in blue; 2.36%± 1.78 for transfer by secretion in green). (**p=0.001 for RNAi Control versus RNAi α-Catenin for N=3). (G, H). Representative confocal images showing 24h co-culture between (G) GFP N-Cadherin SH-SY5Y with DiD-labelled vesicles (donor) and SH-SY5YmCherry cells transfected with RNAi Control (acceptor), (H) co-culture between GFP N-Cadherin challenged with DiD-labelled vesicles (donor) and mCherry cells transfected with RNAi α-Catenin (acceptor). (I, J). Representative confocal images showing 24h co-culture between (I) mCherry cells transfected with RNAi Control with DiD-labelled vesicles (donor) and GFP N-Cadherin cells (acceptor). (J) 24h co-culture between mCherry cells transfected with RNAi α-Catenin challenged with DiD-labelled vesicles (donor) and GFP N-Cadherin cells (acceptor). The yellow arrowheads indicate DiD-labelled vesicles detected in the cytoplasm of acceptor cells. (K). Graph showing the percentage of acceptor cells containing DiD-labelled vesicles from the co-cultures in the conditions described in (G) (44.5%± 7.25 for contact-mediated transfer in blue; 3.04%± 0.80 for transfer by secretion in green), (H) (21.6%± 4.87 for contact-mediated transfer in blue; 2.89%± 0.60 for transfer by secretion in green), (I) (46.21%± 4.29 for contact-mediated transfer in blue; 2.12%± 0.21 for transfer by secretion in green) and (J) (24.37%± 1.29 for contact-mediated transfer in blue; 2.09%± 0.40 for transfer by secretion in green). (***p<0.0001 for (G) versus (H) for N=3; ***p<0.0001 for (I) versus (J) for N=3; ns p=0.9985 for (G) versus (I) for N=3; ns p=0.7643 for (I) versus (J) for N=3). (L). Confocal micrograph showing TNTs between RNAi α-Catenin (mCherry) and N-Cadherin (GFP) cells plated on EM grids. (M). Low cryo-EM micrograph showing TNT-connected cells in in the dashed yellow square in (L). (N). Intermedia cryo-EM micrograph of (M). (O). High-magnification cryo-tomography slices corresponding to the red dashed square in (N). (P) Intermedia cryo-EM micrograph corresponding to the green dashed square in (N). (Q). High-magnification cryo-tomography slices corresponding to the blue arrow in (N). (R). High-magnification cryo-tomography slices corresponding to the pink arrow in (N). (S). Table showing the percentage of TNTs fully extended and closed tip in between RNAi α-Catenin (mCherry) and GFP N-Cadherin. The table is organized by the cell of origin of the iTNTs. Scale bars: (A, B, D, E, G-L) 10 µm, (M, N, P) 2 µm, (O, Q, R) 100nm.

Considering that cadherins are cell adhesion molecules that act in trans, affecting the downstream actin cytoskeleton through α-Catenin, we decided to investigate the effects of trans interaction between these two molecules on the establishment of functional TNTs. To this aim we co-cultured one cell population overexpressing N-Cadherin with another cell population KD for α-Catenin and analyzed both the number of TNTs and their functionality. As for the previous experiments, KD of α-Catenin resulted in 71% of reduction in the levels of this protein compared to the RNAi Control (Fig. S4E), whilst α-Catenin levels in GFP N-Cadherin cells were almost 2-fold compared to the RNAi Control cells (Fig. S4E). In these conditions, there was an increase in both the total percentage of TNT connected cells (Fig. S6A) and in the percentage of heterotypic connections (e.g., the connections between GFP N-Cadherin cells and α-Catenin KD cells) (Fig. S6B) without altering the distribution of connections between the different cell types (Fig. S6C for the RNAi Control and Fig. S6D for the RNAi α-Catenin).

We then performed the DiD-vesicle transfer assay in two different conditions: 1) GFP N-Cadherin cells as donor cells cocultured with either RNAi Control (Fig. 5G) or RNAi α-Catenin cells (Fig 6H) as acceptors; 2) RNAi Control (Fig. 5I) or RNAi α-Catenin cells (Fig. 5J) as donors cocultured with GFP N-Cadherin cells as acceptors. Contact-mediated transfer of DiD-vesicles was around 44% between GFP N-Cadherin cells as donor and RNAi Control cells as acceptors and 46% between RNAi Control cells as donors and GFP N-Cadherin cells as acceptors. Interestingly, KD of α-Catenin either in the donor or in the acceptor population resulted in around 50% of decrease of the contact mediated transfer compared to control conditions with a 22% and 24% of acceptor cells receiving DiD-vesicles respectively for the two aforementioned conditions (Fig. 5K) suggesting that α-Catenin was necessary both in the donor and acceptor cells for a functional TNT to be established. Again, a minimal part of the total transfer corresponded to a secretion mechanism (Fig. 5K). To better understand these results, we analysed the structure of TNTs formed between one cell population overexpressing N-Cadherin and another cell population KD for α-Catenin by correlative cryo-TEM where we could recognize the two cell populations differently labeled in FM (Fig. 5L). As shown in the example of Fig. 5M-R we frequently observed that TNTs established between these two different cell populations corresponded to a bundle of 2 iTNTs (Fig. 5L, Fig. S7). Interestingly, both the iTNT coming from the GFP N-Cadherin cells and the one coming from the RNAi α-Catenin cells had close-ended tips (Fig. 5O-P, Fig. S7, Movie S6). Quantitative analysis of these cryo-EM data revealed that 90% of iTNT originating from GFP N-Cadherin cells and 90% of iTNT originating from RNAi α-Catenin cells were closed-tip iTNTs (Fig. 5S) and only 10% were fully extended between the two cell populations (Fig. 5S). These results strongly indicated α-Catenin working downstream N-Cadherin was needed both in the donor and acceptor cell to establish functional TNTs.

**Figure 6.**
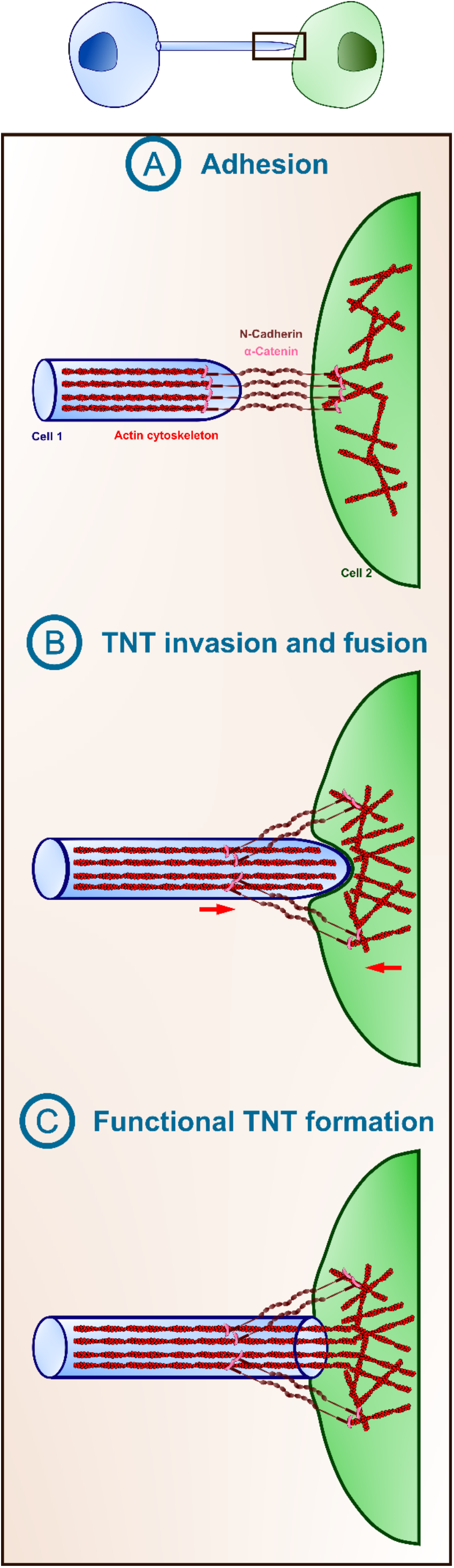
Model of the role of the Cadherin-Catenin complex in the formation of functional TNTs. (A) The tip of the TNTs that is formed from cell 1 would reach the opposing cell 2 and establish direct physical contact adhering through the homophilic interactions of the N-Cadherin/α-Catenin complex, that would anchor both membranes. (B) The TNT would continue protruding towards the opposing cells, producing pushing forces which can result in membrane invagination and pulling and resistance forces (red arrows) in the opposing cell. Similarly to what has been observed in *drosophila* myoblast fusion, this would lead to close proximity of both membranes and the eventual fusion. (C) Once fusion has occurred, a functional TNT it is formed, forming an open channel between both cells that can now exchange cargoes.

## Discussion

Here we show that N-Cadherin interference leads to an increase in TNT-connected cells and a decrease in TNT-mediated vesicle transfer. On the contrary, overexpression of N-Cadherin, results in a decrease in TNT-connected cells but an increase of vesicle transfer. These data uncover a role for N-Cadherin in the functional establishment of TNTs. However, the question arises as to why N-cadherin KD leads to an increase in TNT connected cells while N-Cadherin OE leads to a decrease? Why should there be less transfer in conditions when TNTs increase, and more transfer when TNTs are reduced? Cryo-EM and tomography (23) on RNAi N-Cadherin cells revealed that the canonical structure of the TNTs, i.e. the parallel bundle of iTNTs, was highly altered, with iTNTs crossing over each other and in many cases without specific direction. Conversely, N-Cadherin OE led to an opposite phenotype, with iTNTs highly ordered, running parallel to each other, and directed straight toward the opposing cell. This alteration of the bundle structure of the iTNTs is consistent with a role for N-Cadherin treads in facilitating the organization of the bundle of iTNTs into a highly ordered, parallel and more stable structure. Furthermore, we observed that cells forming TNTs use other pre-existing protrusions as guides to grow (Movie S7 &8), the lack of N-Cadherin would therefore cause the disappearance of these guides and therefore these iTNTs would have no reference for growth/retraction. Our data are supported by recent findings showing that double filopodial bridges (DFB, that the authors consider as precursors of close-ended TNTs in HeLa cells), were dissociated resulting into separation of paired cells by downregulating N-Cadherin (or inhibiting its function with EGTA) (38). In the same study, the authors show N-Cadherin decorating the whole DFB/TNT-like structure and preferentially enriched in the areas of contact with opposite cells. They interpret these enrichments as an indication of close-ended TNTs formation, also supported by the fact that they only observe unidirectional transfer of Ca^2+^ (and not of different cellular material or organelles) in these structures. We found a similar enrichment of N-Cadherin at the TNTs ends (Movie S9 & 10), however, when we overexpress N-Cadherin, in addition to these enrichments we observe a significant increase in vesicle transfer. This, together with our previous ultrastructural study of TNTs (23), suggests that at least in our cellular model these TNTs should be open-ended and that the continuity between TNT and opposite cell seems to be facilitated by N-Cadherin. Indeed, quantitative analysis of our cryo-EM data showed in RNAi N-Cadherin cells there was a substantial increase in the number of close-ended iTNTs in a bundle, thus, explaining the decrease in vesicle transfer. We can therefore speculate that N-Cadherin may also regulate the process of fusion of the TNT with the opposing cell. The involvement of cadherin proteins in cellular fusion has been described in myoblast (42), in trophoblastic cells fusion (43) and in multinucleated osteoclasts formation (44). Nevertheless, N-Cadherin is not able to directly trigger fusion because the distance between two molecules of N-Cadherin on opposite membranes is 37.8 nm (25), which is too big to lead to spontaneous fusion, implying that it could facilitate a pre-fusion event, the adhesion between the opposite cell membranes prior to the fusion. Therefore, in N-Cadherin KD conditions, failure in the adhesion of the TNTs with the opposing cell would impair fusion resulting in close-ended TNTs and reduction of material transfer. Furthermore, in GFP N-Cadherin cells, TNTs were predominantly formed by a single-tube reaching a diameter of 600 nm, and very often containing organelles inside them (Fig. 2H). The presence of larger tubes together with the increase in fully extended connections could further explain the higher transfer observed in conditions of OE of N-Cadherin. One interesting question raised from these data is whether the single larger tubes are derived from iTNTs and what is the role of N-Cadherin in this event. N-Cadherin is present on short linkers between iTNTs, (23; Fig. S2G), it may be possible that the increased presence of N-Cadherin could allow the fusion of iTNTs in single tubes. In conditions of N-Cadherin OE, TNTs were more stable (e.g., lasted longer) compared to KD or control cells. One possible explanation is that the N-Cadherin linkers could stabilize the iTNTs bundles and therefore reduce TNT fragility, as well as, that the single tubes are more robust and last longer compared to the bundles. This is line with recent atomic force microscope (AFM) measurements demonstrating that TNTs are elastic structures and that N-Cadherin regulates flexural strength of the TNTs (45). Further studies will be needed to understand the specificity and nature and origin of iTNTs and single tubes. We observed that α-Catenin KD and OE phenocopied the effects on TNTs of N-Cadherin.Through the link with the actin cytoskeleton, the N-Cadherin and α-Catenin complex might regulate actin dynamics (34, 35), interact with other actin-related proteins such as cortactin (46), Arp2/3 complex (34) or formins (47) and other proteins involved in actin polymerization-depolymerization cycle which is a key step in TNT formation (48). Despite N-Cadherin OE, the lack of α-Catenin is sufficient to recapitulate the observed KD effects of N-Cadherin or α-Catenin in naive cells, showing that α-Catenin is a downstream effector of N-Cadherin in the regulation of TNTs. Finally, we have shown that α-Catenin is necessary in both cell population as KD in donors or acceptors results in decrease in vesicles transfer and increase in closed-tips TNTs. We can speculate that these iTNTs are not functional and are not able to fuse with the opposite cells to share materials. We hypothesize that TNT fusion might resemble myoblast fusion (Fig. 6) (49). In *Drosophila* myoblast fusion, fusion-competent myoblast cells extend F-actin finger-like protrusion that invade the opposing founder cell (50). The membranes of these invasive protrusions and the receiving cell are engaged by cell adhesion molecules (51) that would initiate a signaling cascade towards the cytoskeleton increasing cortical tension by the pushing forces of the protrusions and the pulling of the membrane of the receiving cells that will eventually lead to a pore formation and membrane fusion (52). In our case, we speculate that TNTs protrusions would establish direct physical contact through N-Cadherin/α-Catenin complex with the recipient cell (Fig. 6A), and the tip of the TNT would continue invading the opposing cell, exerting the required push/pull forces that are transmitted to the cortical actin cytoskeleton through α-Catenin (Fig. 6B), forming a fusion competent site so an open-ended connection could be formed (Fig. 6C). Overall, our study begins to shed some light on the mechanisms of formation of such peculiar structure, revealing the essential role and different functions of the N-Cadherin-α-Catenin complex in TNTs in neuronal cells.

## Material and Methods

### Cell lines, plasmids and transfection procedures

Human neuroblastoma (SH-SY5Y) cells were cultured at 37 °C in 5% CO2 in RPMI-1640 (Euroclone), plus 10% fetal bovine serum and 1% penicillin/streptomycin (gift from Simona Paladino, Department of Molecular Medicine and Medical Biotechnology, University of Naples Federico II, Naples, Italy). GFP N-Cadherin plasmid was available in the lab and was obtained from Sandrine Etienne-Manneville (Pasteur Institute, Paris, France) (53, 54), mEmerald α-Catenin plasmid was purchased from Addgene (#53982). To obtain clones that express GFP N-Cadherin or mEmerald α-Catenin, cells were transfected with the corresponding plasmid using Lipofectamine 2000 (Invitrogen) following the manufacture recommendations and selected with 300 ug/mL of geneticin for 10-14 days, changing the medium every 3-4 days. The pool of cells was seeded in 96-well plates through a limiting dilution in such a way that 0.5 cells are seeded per well, and after allowing them to grow, they were analyzed and the clone overexpressing the protein of interest were selected. Human siRNA Oligo Duplex for N-Cadherin (SR300716) and α-Catenin (SR301060) were purchased from Origene. siRNA was transiently transfected to the cells through Lipofectamine RNAimax (Invitrogen) following the manufacture recommendations and the experiments are carried out in between 48 and 72 hours after the transfection.

### Sample preparation for visualization and quantification of the TNTs

SH-SY5Y cells were trypsinized and counted and 100.000 cells were plated overnight (O/N) in coverslips. Cells transfected with the corresponding siRNA were trypsinized and counted at 48 hours post-transfection and 100.000 cells were plated on coverslips O/N. 16 hours later cells were fixed with specific fixatives to preserve TNTs first with fixative solution 1 (2% PFA, 0.05% glutaraldehyde and 0.2 M HEPES in PBS) for 15 min at 37 °C followed by a second fixation for 15 min with fixative solution 2 (4% PFA and 0.2 M HEPES in PBS) at 37 °C (for more information, 40). After fixation cells were washed with PBS and membrane was stained with conjugated Wheat Germ Agglutinin (WGA)-Alexa Fluor (1:300 in PBS) (Invitrogen) and DAPI (1:1000) (Invitrogen) for 15 minutes at room temperature, followed by 3 gentle washes with PBS and finally samples were mounted on glass slides with Aqua PolyMount (Polysciences, Inc.).

### Quantification of TNT-connected cells

Various Z-stacks images of different random points of the samples are acquired with an inverted laser scanning confocal microscope LSM700 (Zeiss) controlled by the Zen software (Zeiss). Images are analyzed following the morphological criteria of the TNTs: structures that connect distant cells and not adherent, so for, first slices are excluded and only connections present in the middle and upper stacks are counted. Cells containing TNTs between them are marked as TNT-connected cells and by counting the number of cells that have TNTs between them and the total number of cells, the percentage of cells connected by TNTs is obtained. Analysis of the TNT-connected cells was performed in ICY software (https://icy.bioimageanalysis.org/) using the “Manual TNT annotation plugin”. At least 200 cells per condition were counted in each experiment. Image were processed with the ImageJ software.

### DiD transfer assay (co-culture assay)

DiD transfer assay is described elsewhere (40) a co-culture is performed consisting of two populations of cells labeled differently: first, your cells of interest (donors) are treated with Vybrant DiD (dialkylcarbocyanines), a lipophilic dye that stains the vesicles, 1:1000 (Thermo Fisher Scientific) in complete medium for 30 minutes at 37 °C (Life Technologies) and second, these cells are co-cultured at a ratio of 1:1 with another population of cells (acceptors) marked in another color (normally cells expressing soluble GFP or soluble mCherry) and grown for about 16 hours. For SH-SY5Y 50.000 donor cells are cocultured with 50.000 acceptor cells on coverslips. The results are analyzed through microscopy as described above and the final results are obtained by semiquantitative analysis with the ICY software from calculating the percentage of acceptor cells with marked vesicles among the total number of acceptor cells. At least 100 acceptor cells per condition were counted in each experiment. Image montages were built afterward in ImageJ software

### Trypsin treatment experiment

Cell singularization by trypsin in was adapted from (41). SH-SY5Y cells were plated the day before the experiment, seeding twice as many cells as under normal conditions, 800.000 cells per condition, since trypsin treatment would cause us to lose part of the cells that would detach in Ibidi µ-dishes (Biovalley, France) to favor cell adhesion with the substrate. 16 hours later the culture medium was replaced by 0.05% Trypsin/EDTA (Gibco), enough to cover the whole dish, for 3 minutes at room temperature. Immediately after these cells were fixed, stained, sealed and analyzed exactly in the same way as described in “sample preparation for visualization and quantification of the TNTs”

### Immunofluorescence

For immunofluorescence, 100.000 cells were seeded on glass coverslips and after O/N culture they were fixed with 4% paraformaldehyde (PFA) for 15 minutes at °C, quenched with 50 mM NH_4_Cl for 15 min, permeabilized with 0.1% Triton X-100 in PBS and blocked in 2% BSA in PBS. Primary antibodies used are: rabbit anti-N-Cadherin (ABCAM ref: ab76057), rabbit anti-N-Cadherin (Genetex ref: GTX127345) mouse anti-N-Cadherin (BD Biosciences ref: 610920), and rabbit anti-α-Catenin (Sigma ref: c2081) all of them at 1:1000 in 2% BSA in PBS during 1 hour. After 3 washes of 10 minutes each with PBS, cells were incubated with each corresponding AlexaFluor-conjugated secondary antibody (Invitrogen) at 1:1000 in 2% BSA in PBS during 1 hour. For those experiments showing the actin cytoskeleton, cells were labeled with Rhodamine Phalloidin (Invitrogen) at 1:1000 in the same mix and conditions as the secondary antibodies. Then, cells were washed 3 times of 10 minutes each with PBS, stained with DAPI and mounted on glass slides with Aqua PolyMount (Polysciences, Inc.). Images were acquired with a confocal microscope LSM700 (Zeiss) and processed with the ImageJ software.

### Western blot

For Western blot cells were lysed with lysis buffer composed by 150 mM NaCl, 20 mM Tris, 5 mM EDTA, pH 8.0. Protein concentration was measured by a Bradford protein assay (Bio-Rad). Samples were boiled at 100 °C for 5 min and loaded in handcrafted 8% SDS-polyacrylamide gel or 4-12% Criterion™ XT Bis-Tris XT Precast Gels (Bio-Rad) and electrophoresed in 1X Tris/Glycine/SDS buffer (Bio-Rad) or 1X XT MOPS buffer (Bio-Rad) respectively for 1.5-2 hours at 90V. Proteins were transferred to 0.45 µm Nitrocellulose membranes (Bio-Rad) with 1X Tris/Glycine transfer buffer (Bio-Rad) for 1.5 hours at 90V in a cold chamber. Membranes were blocked in 5% non-fat milk in Tris-buffered saline with 0.1% Tween 20 (TBS-T) for 1 hour. Membranes were incubated O/N at 4 °C with the corresponding primary antibodies at 1:1000 in 5% non-fat milk TBS-T. Primary antibodies used for Western blot were: rabbit anti-N-Cadherin (ABCAM ref: ab76057), rabbit anti-N-Cadherin (Genetex ref: GTX127345) mouse anti-N-Cadherin (BD Biosciences ref: 610920), rabbit anti-α-Catenin (Sigma ref: c2081) and mouse anti-α-tubulin (Sigma ref: T9026). Membranes were washed 3 times 10 minutes each with TBS-T and then incubated with the corresponding IgG secondary antibodies horseradish peroxidase-conjugated (GE Healthcare Life Sciences) at 1:1000 for 1 hour at room temperature. Membranes were washed 3 times 10 minutes each. Membrane protein bands were detected with Amersham™ ECL Prime Western Blotting Detection Reagent (Cytiva). Membranes were imaged using Amersham™ Imager 680 (GE Healthcare Life Sciences).

### Live Imaging

400.000 SH-SY5Y cells were plated the day before the experiment in Ibidi µ-dishes. After 16 hours of culture, live time series images were acquired with a 60 × 1.4NA CSU oil immersion objective lens on an inverted Elipse Ti microscope system (Nikon Instruments, Melville, NY, USA). Cells were labeled with 1:1000 dilution of conjugated WGA-Alexa Fluor in the corresponding media. Images were captured in immediate succession with one of two cameras, which enabled time intervals between 20 and 40 seconds per z-stack or between 50 and 70 seconds per z-stack when using two lasers. For live cell imaging, the 37 °C temperature was controlled with an Air Stream Stage Incubator, which also controlled humidity. Cells were incubated with 5% CO2 during image acquisition.

### Cell preparation for cryo-EM

Carbon-coated gold TEM grids (Quantifoil NH2A R2/2) were glow-discharged at 2 mA and 1.5– × 10^−1^ m bar for 1 minute in an ELMO (Cordouan) glow discharge system. Grids were sterilized under UV three times for 30 minutes at R. T. and then incubated at 37 °C in complete culture medium for 2 hours. 300,000 SH-SY5Y cells (RNAi N-Cadherin/α-Catenin, GFP N-Cadherin/mEmerald α-Catenin) were counted and seed on cryo-EM grids positioned in 35 mm Ibidi µ-Dish (Biovalley, France). After 24 hours of incubation, resulted in 3 to 4 cells per grid square. Prior to chemical and cryo-plunging freezing, cells were labeled with WGA (1:300 in PBS) for 5 min at 37 °C. For correlative light- and cryo-electron microscopy, cells were chemically fixed in 2% PFA + 0.05% GA in 0.2 M Hepes for 15 minutes followed by fixation in 4% PFA in 0.2 M Hepes for 15 minutes and kept hydrated in PBS-1X buffer prior to vitrification.

For cell vitrification, cells were blotted from the back side of the grid for 10 seconds and rapidly frozen in liquid ethane using a Leica EMGP system as we performed before (16).

### Cryo-electron tomography data acquisition and tomogram reconstruction

The cryo-EM data was collected from different grids at the Nanoimaging core facility of the Institut Pasteur using a Thermo Scientific (TF) 300kV Titan Krios G3 cryo-transmission electron microscopes (Cryo-TEM) equipped with a Gatan energy filter bioquantum/K3. Tomography software from Thermo Scientific was used to acquire the data. Tomograms were acquired using dose-symmetric tilt scheme, a +/-60 degree tilt range with a tilt step 2 was used to acquire the tilt series. Tilt images were acquired in counting mode with a calibrated physical pixel size of 3.2 Å and total dose over the full tilt series of 3.295 e- /Å2 and dose rate of 39,739 e-/px/s with an exposure time of 1s. The defocus applied was in a range of -3 to – 6 µm defocus.

Cryo-EM and tomography (Fig. 1 and S1) was performed on a Tecnai 20 equipped with a field emission gun and operated at 200 kV (Thermo Fisher company). Images were recorded using SerialEM software on a Falcon II (FEI, Thermo Fisher) direct electron detector, with a 14 µm pixel size. Tilt series of TNTs were acquired covering either an angular range of – 52° to + 52°. The defocuses used were −6 µm.

The tomograms were reconstructed using IMOD (eTomo). Final alignments were done by using 10 nm fiducial gold particles coated with BSA (BSA Gold Tracer, EMS). Gold beads were manually selected and automatically tracked. The fiducial model was corrected in all cases where the automatic tracking failed. Tomograms were binned 2x corresponding to a pixel size of 0.676 nm for the Titan and SIRT-like filter option in eTomo was applied. For visualization purposes, the reconstructed volumes were processed by a Gaussian filter.

Quantitative manual measurements of iTNTs full extended, tip-closed, single TNTs were performed considering 10 cryo-EM slices and/or tomograms for control cells, 18 cryo-EM slices and/or tomograms for RNAi N-Cadherin, 45 for GFP N-Cadherin, 16 cryo-EM slices and/or tomograms for RNAi α-Catenin, 18 cryo-EM slices and/or tomograms for mEmerald α-Catenin, 10 cryo-EM slices and/or tomograms for co-culture of GFP N-Cadherin and RNAi α-Catenin.

### Cryo-EM N-Cadherin immuno-labeling

SH-SY5Y cells were plated on grids as described in above. After incubation O/N at 37 °C, cells were fixed with PFA 4% for 20 min at 37 °C, quenched with 50 mM NH4Cl for 15 min, and blocked with PBS containing 2% BSA (w/v) for 30 min at 37 °C. Cells were labeled with a rabbit anti-N-Cadherin ABCAM 76057 antibody (1:200), followed by Protein A-gold conjugated to 10 nm colloidal gold particles (CMC, Utrecht, Netherlands). SH-SY5Y cells were then rapidly frozen in liquid ethane as above.

### Statistical analysis

The statistical analysis for the experiments concerning the percentage of TNT-connected cells and the DiD transfer assay are described elsewhere (6). Briefly, the statistical tests were computed using either a logistic regression model computed using the ‘glm’ function of R software (https://www.R-project.org/) or a mixed effect logistic regression model using the lmer and lmerTest R packages, applying a pairwise comparison test. For the rest of experiments, Student’s t-test (for 2 groups) or One-Way ANOVA (for more than 2 groups) tests were applied. All column graphs, Student’s t-test and One-Way ANOVA statistical analysis were performed using GraphPad Prism version 9 software.

## Acknowledgments

We thank all the laboratory members for useful discussion. We are grateful to R. Bouyssie, a member of the administrative staff of the Membrane Traffic and Pathogenesis department at Institut Pasteur. The NanoImaging Core at Institut Pasteur is acknowledged for support with image acquisition and analysis, particularly M. Vos, and J.-M. Winter (NanoImaging Core at Institut Pasteur). We also acknowledge G. Péhau-Arnaudet (Ultrapole, Institut Pasteur).

## Author contributions

A.P., R.N.M., and C.Z. conceived the experiments. A.P. and R.N.M. prepared the figures and wrote the manuscript; R.N.M. performed cocultures, TNT, and WB quantifications. A.P. set up and performed all correlative, cryo-CLEM and cryo-ET experiments by using TITAN cryo-EM, tomograms reconstruction and quantitative analysis. A.S. performed EM acquisition by using Falcon F20. C. B. discussed experiments. A.P., R.N.M., C.B., and C.Z. discussed the results. All authors commented on the manuscript. C.Z. conceived the project, supervised all the work, and wrote the manuscript. C.Z. contributed to funding acquisition.

