## Supplementary Figures for "N-Cadherin and α-catenin regulate formation of functional tunneling nanotubes"

### Supplementary Materials

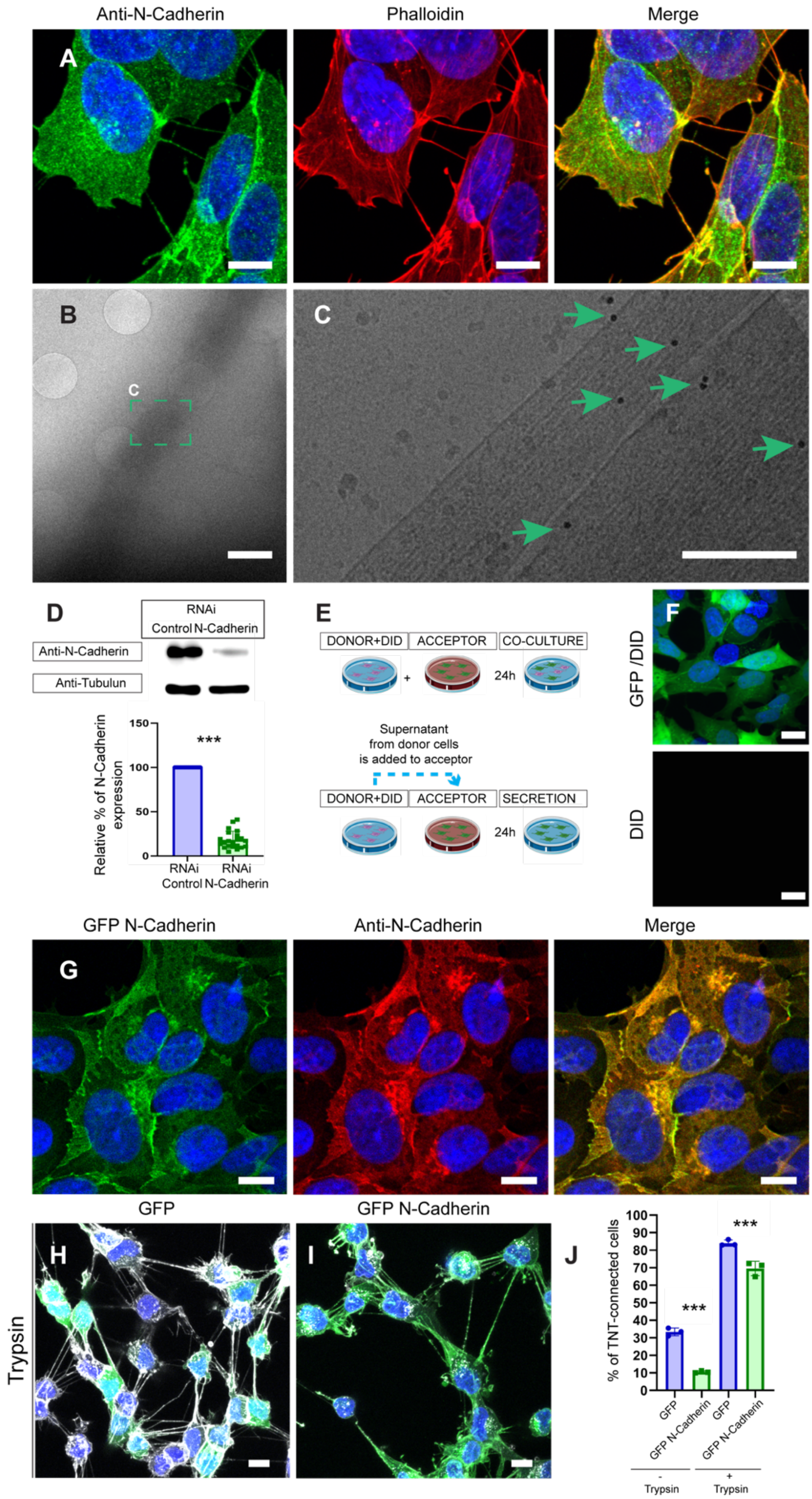

**Fig. S1. Confocal and ultrastructure analysis of N-Cadherin in TNT connected SH-SY5Y cells and co-culture and secretion pipeline.**

**(A).** Confocal micrograph of the immunofluorescence of N-Cadherin showing TNT connected SH-SY5Y cells. N-Cadherin (green) and actin (red). Cells stained and DAPI (blue) for the nuclei. **(B, C).** Immunogold EM anti-N-Cadherin (B) low magnification micrograph showing a TNT connecting SH-SY5Y cells, (C) high magnification of green dashed squares in (B) showing the presence of N-Cadherin (green arrows) on iTNTs. **(D).** Western blot of RNAi Control cells and RNAi N-Cadherin. Membrane was blotted with antibodies anti N-Cadherin and α-tubulin as loading control (top). Graph showing the relative expression of N-Cadherin in RNAi Control cells (100%) and RNAi N-Cadherin (18.1% ± 9.81) (***$p<0.0001$ for RNAi Control versus RNAi N-Cadherin for N=22) (bottom). **(E).** (top) Description of co-culture experiments: Donor cells stained with DiD labeled vesicles cells were co-cultured with the acceptor cells and an incubate for additional 24h before to be fixed. (bottom) Description of secretion experiments: the medium from DiD-stained donor cells was added on acceptor cells for 24h. **(F).** Representative confocal micrograph showing acceptor cells (GFP-labeled) that have received supernatant from DiD-labeled donor cells. Cells stained with DAPI (blue) for the nuclei. **(G).** Immunofluorescence of N-Cadherin (red) in GFP N-Cadherin cells. Cells stained with DAPI (blue) for the nuclei. **(H, I).** Confocal micrograph showing (E) TNTs between GFP cells treated with Trypsin-EDTA, (F) TNTs between GFP N-Cadherin cells treated with Trypsin-EDTA. **(J).** Graph showing the percentage of TNT-connected cells in GFP cells (84.1% ± 1.80) and GFP N-Cadherin cells (69.6% ± 4.13) treated with Trypsin-EDTA (***$p<0.0001$ for GFP versus GFP N-Cadherin for N=3). Cells stained with WGA-647 (gray) and DAPI (blue) for the nuclei. Scale bars in **(A, G, H, I)** 10 μm, **(B)** 2 μm, **(C)** 100nm, **(F)** 20 μm.

RNAi N-Cadherin

GFP N-Cadherin

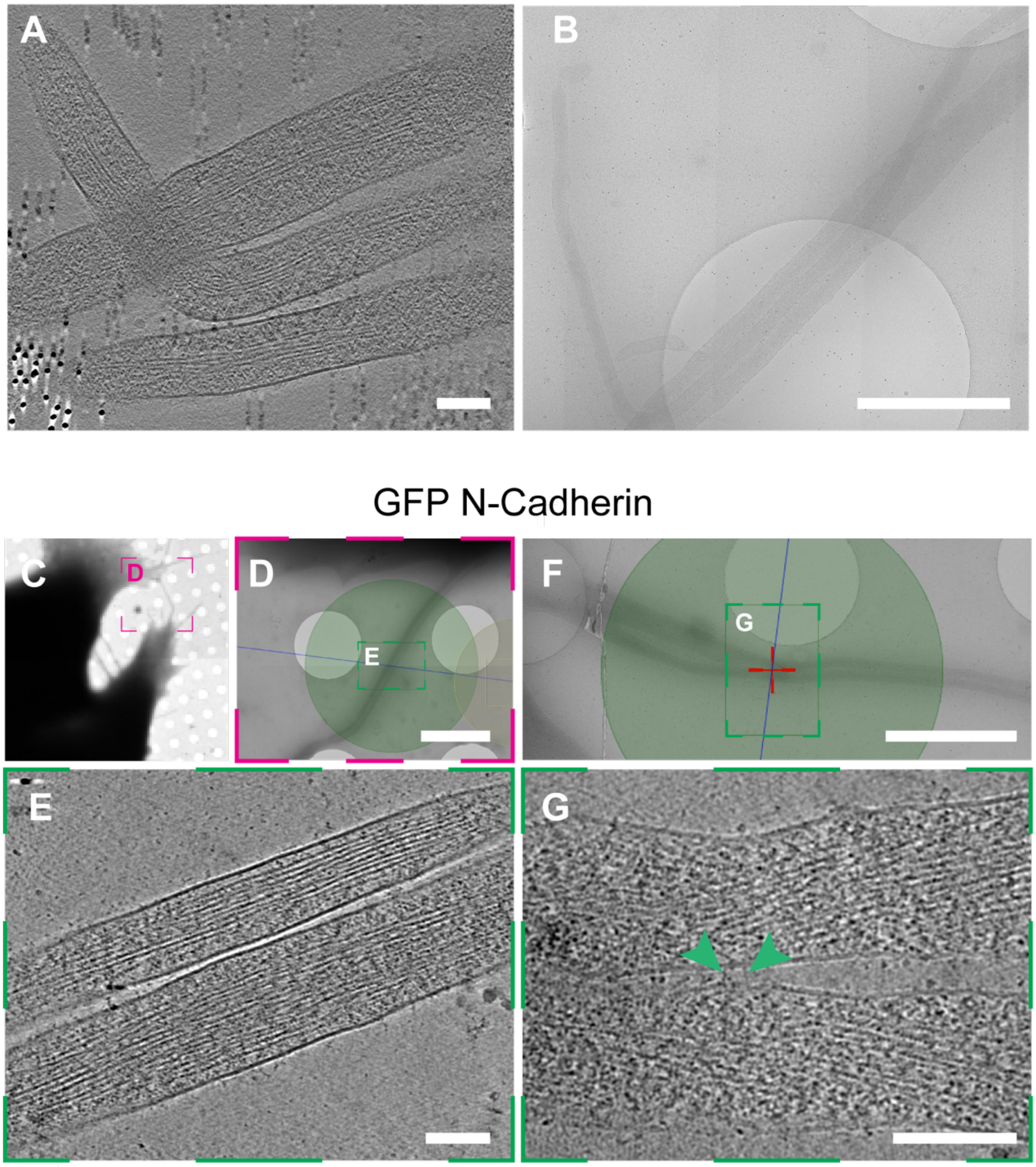


H

| | Ultrastructure iTNTs | |
|---|---|---|
| | % fully extended | % closed tip |
| **GFP N-Cadherin cells** | 58% | 42% |

**Fig. S2. Cryo-EM on TNTs formed between SH-SY5Y in which N-Cadherin was up or down-regulated.**

**(A, B).** Cryo-EM grids were prepared using RNAi N-Cadherin cells (A) High-magnification cryo-tomography slices showing iTNTs with closed tip (B) intermedia magnification of cryo-EM micrograph showing iTNTs connecting RNAi N-Cadherin cells with closed tip and iTNTs did not run parallel and braided over each other. **(C-G).** Cryo-EM grids were prepared using GFP N-Cadherin cells. (C) Low (D) and intermedia (E) magnification of cryo-EM micrograph showing TNT GFP N-Cadherin cells. (F) Intermediate magnification of cryo-EM micrograph showing TNT connecting GFP N-Cadherin cells. (G) High-magnification cryo-tomography slices of the green dashed square in (F). **(H).** Table showing the percentage of TNTs full extended and closed tip in GFP N-Cadherin cells. Scale bars **(A, E, G)** 100nm, **(B, F)** 2µm.

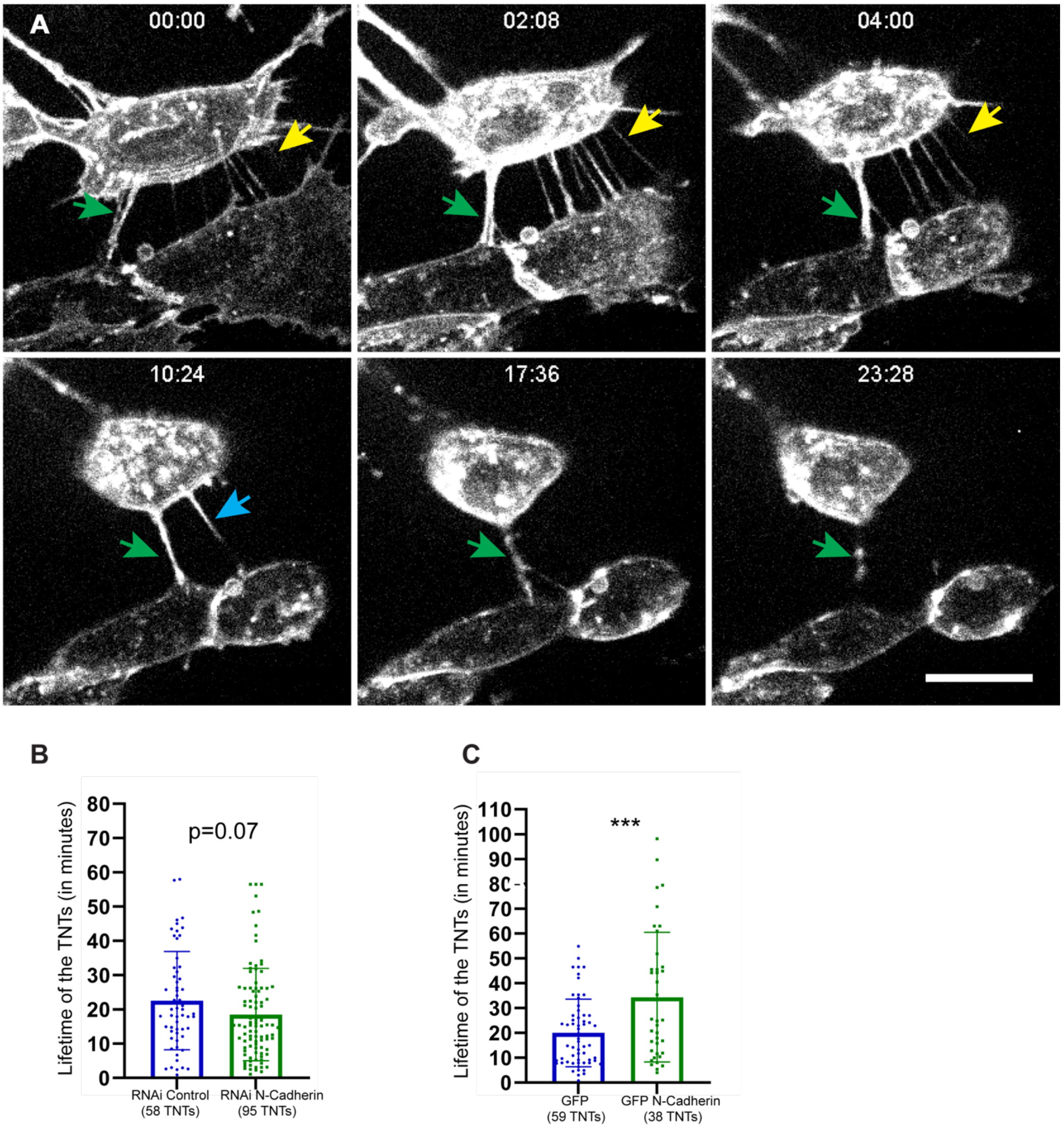


**Fig. S3. N-Cadherin modulates the stability of the TNTs.** (**A**). Representative snapshots of TNTs over time from Movie S2. Arrows point to the TNTs. (**B**). TNT's average duration in RNAi Control or RNAi N-Cadherin cells. Cells were stained with WGA-488 to visualize the cell membrane and TNTs. The graph represents the average lifetime ± SD of 58 TNTs in RNAi Control (22.5 minutes ± 14.33) and 95 TNTs in RNAi N-Cadherin (18.5 minutes ± 13.44) (ns p=0.0795 for RNAi Control versus RNAi N-Cadherin). (**C**). TNT's average duration in control GFP cells or overexpressing N-cadherin. Cells were stained with WGA-647 to visualize the cell membrane and TNTs. Graph represents the average lifetime ± SD of 59 TNTs in GFP control cells (20 minutes ± 13.63) and 38 TNTs in GFP N-Cadherin cells (34.3 minutes ± 26.06) (***p=0.0006 for GFP versus GFP N-Cadherin). Scale bars in (A) correspond to 10 μm.

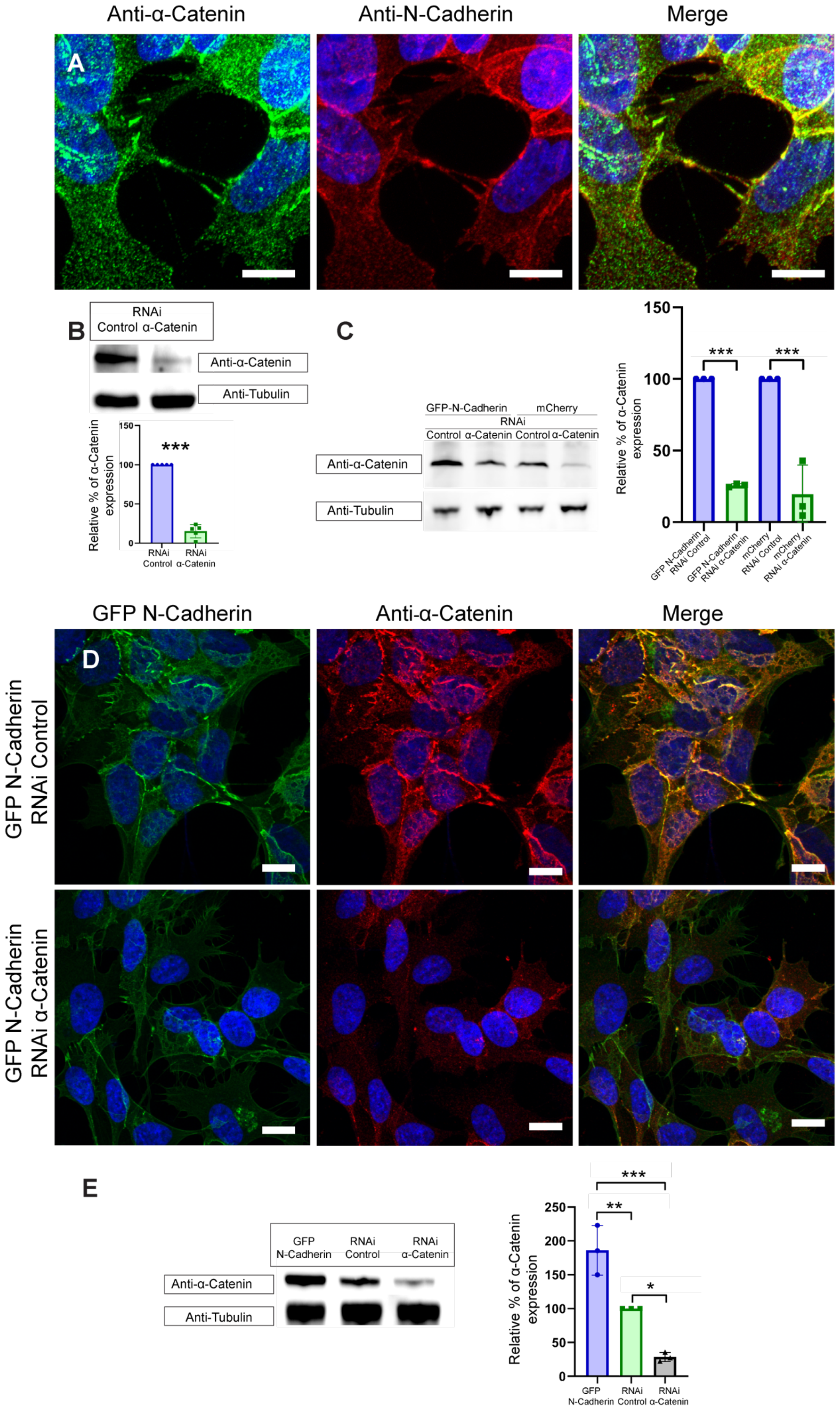
Anti-α-Catenin
Anti-N-Cadherin
Merge
A
B
RNAi
Control α-Catenin
Anti-α-Catenin
Anti-Tubulin
Relative % of α-Catenin expression
150
100
50
0
***
RNAi Control
RNAi α-Catenin
C
GFP-N-Cadherin
mCherry
RNAi
Control α-Catenin Control α-Catenin
Anti-α-Catenin
Anti-Tubulin
Relative % of α-Catenin expression
150
100
50
0
***
***
GFP N-Cadherin RNAi Control
GFP N-Cadherin RNAi α-Catenin
mCherry RNAi Control
mCherry RNAi α-Catenin
GFP N-Cadherin
Anti-α-Catenin
Merge
D
GFP N-Cadherin RNAi Control
GFP N-Cadherin RNAi α-Catenin
E
GFP N-Cadherin
RNAi Control
RNAi α-Catenin
Anti-α-Catenin
Anti-Tubulin
Relative % of α-Catenin expression
250
200
150
100
50
0
***
**
*
GFP N-Cadherin
RNAi Control
RNAi α-Catenin

**Fig. S4. Expression of α-Catenin in RNAi α-Catenin and RNAi N-Cadherin cells and N-Cadherin OE and α-Catenin KD.**

**(A).** Confocal micrograph showing α-Catenin (green) and N-Cadherin (red) IF in cells connected through TNTs. DAPI (blue) for the nuclei. **(B).** Western blot of RNAi Control and RNAi α-Catenin cells. Membrane was blotted with antibodies anti α-Catenin and α-tubulin. Graph shows the relative expression of α-Catenin in RNAi Control cells (100%) and RNAi α-Catenin (15.3% ± 8.49) of all the experiments concerning the knock-down of α-Catenin (***p<0.0001 for RNAi Control versus RNAi α-Catenin for N=5). **(C).** Western blot of RNAi Control and RNAi α-Catenin cells in GFP N-Cadherin cells and mCherry cells. Membrane was blotted with anti α-Catenin and α-tubulin antibodies. Graph shows the relative expression of α-Catenin in GFP N-Cadherin RNAi Control cells (100%) and GFP N-Cadherin RNAi α-Catenin (25.65% ± 1.16), and mCherry RNAi Control cells (100%) and mCherry RNAi α-Catenin (19.51% ± 20.47) (***p<0.0001 for RNAi Control versus RNAi α-Catenin for N=3 in GFP N-Cadherin and mCherry cells). **(D).** Confocal micrograph of α-Catenin (red) IF in GFP N-Cadherin RNAi Control cells and GFP N-Cadherin RNAi α-Catenin cells. Cells stained with DAPI (blue) for the nuclei. **(E).** Western blot of RNAi Control, RNAi α-Catenin and GFP N-Cadherin cells. Membrane was blotted with antibodies anti α-Catenin and α-tubulin. Graph shows the relative expression of α-Catenin in RNAi Control cells (100%), RNAi α-Catenin (28.6% ± 6.75) and GFP N-Cadherin cells (186.1% ± 36.62) (*p=0.0155 for RNAi Control versus RNAi α-Catenin for N=3, **p=0.0065 for RNAi Control versus GFP N-Cadherin for N=3, **p=0.0003 for RNAi α-Catenin GFP N-Cadherin versus for N=3). Scale bars 10 μm.

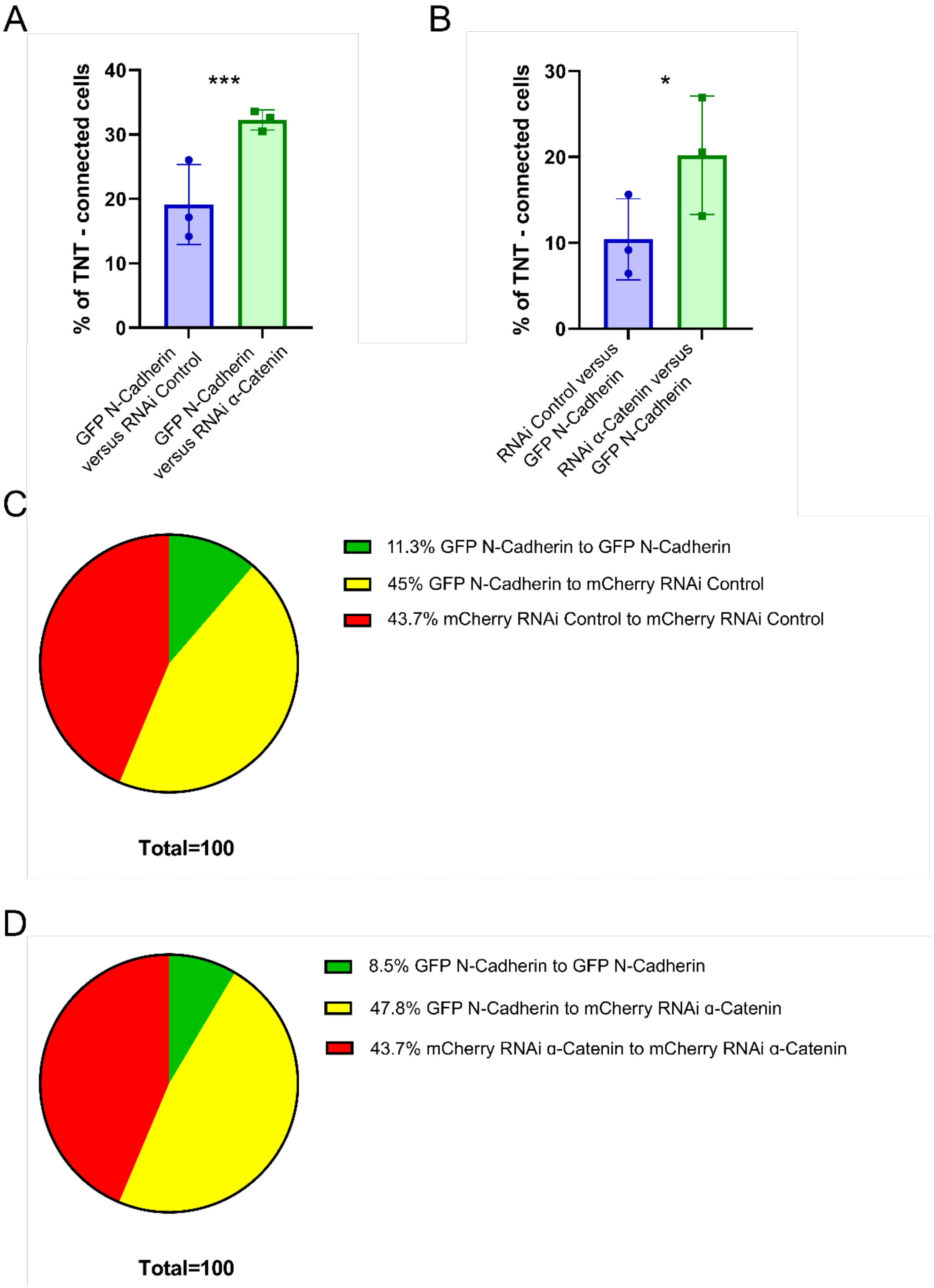


**Fig. S5. % and distribution of TNT connections in the co-culture of cells OE N-Cadherin versus cells interfered for α-Catenin.**

**(A).** Graph showing the total percentage of TNT-connected cells from the co-cultures in GFP N-Cadherin cells versus cells transfected with RNAi Control (19.1%± 6.19) or RNAi α-Catenin (32.2%± 1.58). (***p=0.0004 for RNAi Control versus RNAi α-Catenin for N=3). **(B).** Graph showing the percentage of TNT-connected cells by heterotypic connections (connections made by GFP N-Cadherin cells with RNAi Control or RNAi α-Catenin) from the co-cultures in GFP N-Cadherin cells versus cells transfected with RNAi Control (10.4%± 4.73) or RNAi α-Catenin (20.2%± 6.91). (*p=0.0143 for RNAi Control versus RNAi α-Catenin for N=3). **(C).** Graph showing the distribution of TNTs in the co-culture between GFP N-Cadherin cells and cells transfected with RNAi Control (more than 300 cells counting in each experiment; N =3). Out of the total cells connected, 11.3% corresponds to GFP N-Cadherin cells connected to GFP N-Cadherin cells, 45% corresponds to GFP N-Cadherin cells connected to mCherry RNAi Control cells and 43.7% corresponds to mCherry RNAi Control cells connected to mCherry RNAi Control cells. **(D).** Graph showing the distribution of the connections in the co-culture of GFP N-Cadherin cells versus cells transfected with RNAi α-Catenin (more than 300 cells counting in each experiment; N =3). Out of the total connected TNT cells, 8.5% corresponds to GFP N-Cadherin cells connected to GFP N-Cadherin cells, 47.8% corresponds to GFP N-Cadherin cells connected to mCherry RNAi α-Catenin cells and 43.7% corresponds to mCherry RNAi α-Catenin cells connected to mCherry RNAi α-Catenin cells.

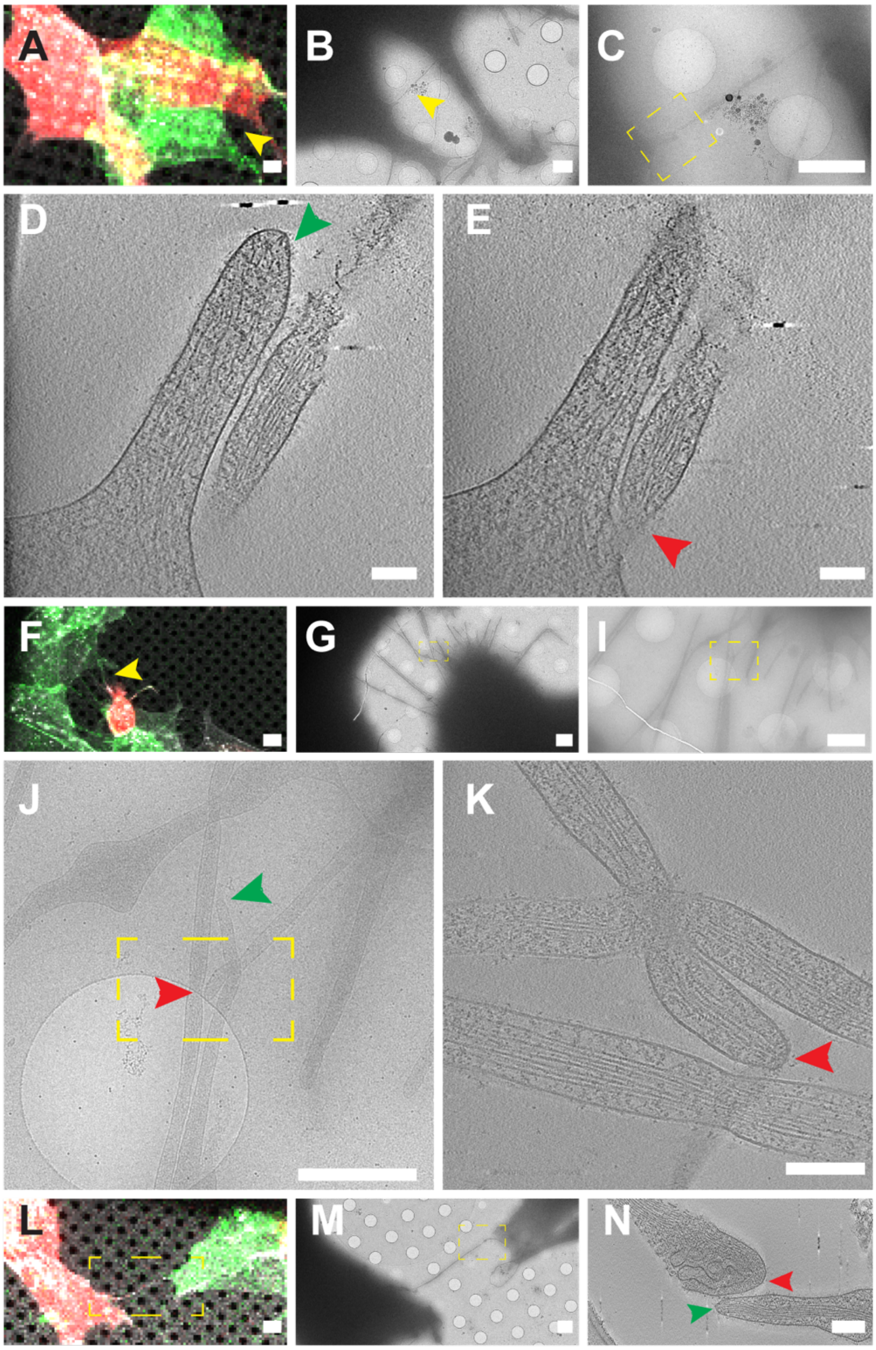


**Fig. S6. Gallery of ultrastructure of TNTs between GFP N-Cadherin and α-Catenin RNAi cells in co-culture.**

**(A, F, L).** Confocal micrograph showing TNTs between RNAi α-Catenin (mCherry) and N-Cadherin (GFP) cells plated on EM grids. **(B, G).** Low cryo-EM micrograph showing TNT-connected cells in (A), (F), indicated by the yellow arrowhead and TNT in the dashed yellow square in (L). **(C, I, M).** Intermedia cryo-EM micrograph of (B), (G), (L) respectively. **(J).** Intermedia cryo-EM micrograph of the yellow dashed square in (I). **(D, E).** High-magnification cryo-tomography slices corresponding to the yellow dashed square in (C), red arrowhead indicates TNT coming from KD α-Catenin cells and green arrowhead indicates TNT coming from GFP N-Cadherin. **(K, N).** High-magnification cryo-tomography slices corresponding to the yellow dashed square in (J) and (M) respectively, red arrowhead indicates TNT coming from KD α-Catenin cells and green arrowhead indicates TNT coming from GFP N-Cadherin. Scale bars in **(A, F, L)** 10 μm, **(B, C, G, I, J, M)** 2 μm, **(D, E, K, N)** 100nm.

**Description of Supplementary Files**

**File Name: Movie S1**

**Description:** Representative slices of a reconstructed tomogram displaying fully extended iTNT connecting two GFP N-Cadherin SH-SY5Y cells shown in Figure S2G. Scale bar: 100nm.

**File Name: Movie S2**

**Description:** Stability measurements in already formed TNTs. Example of a time lapse video of already formed TNTs in which their persistency was measured. Cells were stained with WGA to visualize the membrane (grey). Time between steps: 16 seconds. Scale bar 10 µm.

**File Name: Movie S3**

**Description:** Measurement of the stability of the TNTs in RNAi Control cells. Example of a time lapse video of TNT duration in RNAi Control cells. Cells were stained with WGA to visualize the membrane (grey). Time between steps: 19 seconds. Scale bar 10 µm.

**File Name: Movie S4**

**Description:** Measurement of the stability of the TNTs in RNAi N-Cadherin cells. Example of a time lapse video of TNT duration in RNAi N-Cadherin cells. Cells were stained with WGA to visualize the membrane (grey). Time between steps: 15 seconds. Scale bar 10 µm.

**File Name: Movie S5**

**Description:** Measurement of the stability of the TNTs in GFP N-Cadherin cells. Example of a time lapse video of TNT duration in GFP N-Cadherin cells. Cells were stained with WGA to visualize the membrane. WGA signal it is shown on the left panel and GFP N-Cadherin signal it is shown on the right (both in grey). Time between steps: 52 seconds. Scale bar 10 µm.

**File Name: Movie S6**

**Description:** Representative slices of a reconstructed tomogram displaying iTNT closed tips connecting between GFP N-Cadherin and α-Catenin RNAi cells in co-culture shown in Figure 7D. Scale bar: 100nm.

**File Name: Movie S7**

**Description:** TNT formed *de novo* on top of another TNT in WT cells. Example of a time lapse video of a TNT using a preexisting TNT as a guide to grow towards an opposite cell. Cells were stained with WGA to visualize the membrane (grey). Time between steps: 16 seconds. Scale bar 10 µm.

**File Name: Movie S8**

**Description: TNT formed *de novo* by two dorsal filopodia in WT cells.** Example of a time lapse video of a TNT formed by the interaction on two dorsal filopodia similar to the formation of "Double Filopodial Bridges". Cells were stained with WGA to visualize the membrane (grey). Time between steps: 20 seconds. Scale bar 10 µm.

**File Name: Movie S9**

**Description:** TNT formed *de novo* by cell dislodgement mechanism in GFP N-Cadherin cells. Example of a time lapse video of TNTs formed by the separation of the cell bodies of two cells in GFP N-Cadherin cells and showing accumulation of GFP N-Cadherin signal at the end of these TNTs. Grey signal corresponds to GFP N-Cadherin. Time between steps: 19 seconds. Scale bar 10 µm.

**File Name: Movie S10**

**Description:** Formation of TNTs and accumulation of N-Cadherin at the end of TNTs in GFP N-Cadherin cells. Example of a time lapse video of TNTs formed by cell dislodgement mechanism in GFP N-Cadherin cells and accumulation of GFP N-Cadherin signal at the tips of these TNTs. Grey signal corresponds to GFP N-Cadherin. Time between steps: 48 seconds. Scale bar 10 µm.
